# Regulation of Na_v_1.6-mediated sodium currents underlie the homeostatic control of neuronal intrinsic excitability in the optic tectum of the developing *Xenopus laevis* tadpole

**DOI:** 10.1101/2021.10.07.463558

**Authors:** Adrian C. Thompson, Carlos D. Aizenman

## Abstract

For individual neurons to function appropriately within a network that is undergoing synaptic reorganization and refinement due to developmental or experience-dependent changes in circuit activity, they must homeostatically adapt their intrinsic excitability to maintain a consistent output despite the changing levels of synaptic input. This homeostatic plasticity of excitability is particularly important for the development of sensory circuits, where subtle deficits in neuronal and circuit function cause developmental disorders including autism spectrum disorder and epilepsy. Despite the critical importance of this process for normal circuit development, the molecular mechanism by which this homeostatic control of intrinsic excitability is regulated is not fully understood. Here, we demonstrate that *Xenopus* optic tectal neurons express distinct fast, persistent and resurgent Na^+^ currents. Here, we demonstrate that *Xenopus* optic tectal neurons express distinct fast, persistent and resurgent Na^+^ currents. These are regulated with developmental changes in synaptic input, and homeostatically in response to changes in visual input. We show that expression of the voltage-gated Na^+^ channel subtype Na_v_1.6 is regulated with changes in intrinsic excitability, that blocking Na_v_1.6 channels is sufficient to decrease intrinsic excitability. Furthermore, that upregulation of Na_v_1.6 expression is necessary for experience-dependent increases in Na^+^ currents and intrinsic excitability. Finally, by examining behaviors that rely on visual and multisensory integration, we extend these findings to show that tight regulation of Na^+^ channel gene expression during a critical period of tectal circuit development is required for the normal functional development of the tectal circuitry.

## INTRODUCTION

A key feature of developing circuits is their ability to build highly specialized networks that can process and relay information even while the circuit continues to develop. Neurons of the developing nervous system face a difficult task. They must remain flexible so that they can respond to massive changes in circuit and synaptic organization that occur with development or in response to experience, while maintaining the ability to robustly respond to input by generating a consistent response (Marder and Goaillard, 2006; Turrigiano, 2012; Tien and Kerschensteiner, 2018). To achieve this, neurons can adapt the number, size and strength of synaptic inputs (Turrigiano, 2008). They can also adapt firing rates in response to changing synaptic input (Schulz, 2006). Intrinsic excitability, being the propensity of a neuron to fire an action potential in response to an input signal, is primarily dictated by the distribution and function of voltage-gated ion channels (Raman et al., 1997; Rush et al., 2005). In particular, voltage-gated sodium (Na^+^) channels are strategically placed to mediate the plasticity of neuronal excitability because of their influence on the action potential waveform and threshold (Blair and Bean, 2002; Van Wart and Matthews, 2006). Indeed, changes in Na^+^ current amplitude and kinetics have been shown to modulate excitability (Raman et al., 1997; Rush et al., 2007; Khaliq and Bean, 2010). However, the molecular determinates involved in dynamic changes in Na^+^ currents to regulate homeostatic changes in intrinsic excitability remain elusive.

*Xenopus laevis* tadpoles perform visually-guided behaviours even as the optic tectum, the principal midbrain structure involved sensory integration, continues to develop. Thus, making them an ideal model to study cellular mechanisms by which neurons adapt their intrinsic excitability within the context of a developing, functional neural circuit. The *Xenopus* optic tectum relies on the tight regulation of intrinsic excitability for normal development of this sensory circuit (Dong and Aizenman, 2012), with prior work having shown that homeostatic changes in intrinsic excitability occur primarily with changes in the amplitude of fast Na^+^ current in response to changing levels of excitatory synaptic transmission (Aizenman et al., 2003; Pratt and Aizenman, 2007; Ciarleglio et al., 2015). However, little is known about how changes in Na^+^ currents actually occur, nor of the molecular underpinnings of voltage gated currents in this system. Knowledge of these mechanisms can provide important insights as to how neurons regulate excitability to ensure that they continue to function correctly even as the wider circuit continues to develop and undergo structural and functional rearrangement.

Voltage-gated Na^+^ channels mediate distinct fast, persistent and resurgent Na^+^ currents that display characteristic time scales, voltage-dependencies, and gating properties; all of which can influence neuronal excitability. In voltage-clamp, the short-lived large Na^+^ current evoked by a step depolarization is called the transient Na^+^ current, whereas the relatively smaller, residual current that endures throughout the step is called the persistent Na^+^ current (Hodgkin and Huxley, 1952; French, 1990; Baker and Bostock, 1997). Persistent Na^+^ currents can promote input integration by amplifying dendritic excitatory postsynaptic potentials (Stuart and Sakmann, 1994; Fricker and Miles, 2000), mediating depolarizing afterpotentials (Yue et al., 2005; Chen et al., 2011; Ceballos et al., 2017), as well as regulating neuronal excitability (Wu et al., 2005; Kole, 2011). Additionally, in many cell types, non-conducting voltage-gated Na^+^ channels can reopen in response to a repolarizing step that follows an initial depolarizing step, with the resulting current termed a resurgent Na^+^ current (Cannon and Bean, 2010; Lewis and Raman, 2014). This resurgent Na^+^ current is distinct from persistent Na^+^ current because it is dynamically gated - with distinct rising, peak and falling phases (Raman and Bean, 2001; Bant and Raman, 2010). By generating regenerative depolarizing currents in the voltage range between resting membrane potential and threshold potential, persistent and resurgent sodium currents can play a major role in determining the firing pattern of neurons (Raman et al., 1997; Cummins et al., 2005; Bean, 2007; Ghitani et al., 2016). Fast and persistent Na^+^ currents have been identified in *Xenopus* tectal neurons (Aizenman et al., 2003; Hamodi and Pratt, 2014), with fast Na^+^ currents known to be regulated with changes in neuronal excitability across development and with experience-dependent changes in synaptic strength (Aizenman et al., 2003; Pratt and Aizenman, 2007; Hamodi and Pratt, 2014). It is not known, however, whether resurgent currents exist in this developing system or whether they can be regulated by experience. We also do not understand the molecular mechanisms by which Na^+^ currents are regulated with homeostatic changes in neuronal excitability.

In the present study, we explore the relationship between voltage-gated Na^+^ and K^+^ currents and the regulation of intrinsic excitability in tectal neurons. We measure the association between amplitude of distinct Na^+^ currents and intrinsic excitability in tectal neurons, showing how fast, persistent and resurgent sodium currents are dynamically regulated with changes in intrinsic excitability associated with development and short-term (4hr) enhanced visual stimulation (EVS). We next show how inhibition of Na_v_1.6 channels modulates the resurgent sodium currents and intrinsic excitability of tectal neurons, and illustrate the requirement of Na_v_1.6 channels for the homeostatic increase in sodium currents and intrinsic excitability in response to EVS. Finally, we demonstrate a mechanistic link between Na_v_1.6 channels-mediated increases in intrinsic excitability during a critical period of development and the functional development of tectal circuitry using behavioral assays that test visual and multisensory integration by the optic tectum. These findings illustrate the vital importance of Na_v_1.6 channel expression and distinct Na^+^ currents for the homeostatic control of neuronal excitability in the developing nervous system, and for the first time show that regulation of Na^+^ channel expression is an important mechanism for intrinsic plasticity in the midbrain.

## RESULTS

### The intrinsic excitability of *Xenopus* tectal neurons is correlated with the amplitude of voltage-gated sodium currents

Intrinsic excitability is homeostatically regulated to ensure neurons stably respond to variable synaptic inputs, even when circuits are undergoing dramatic reorganization, such as occurs during development or in response to a changing sensory environment (Pratt and Aizenman, 2007). The intrinsic excitability of tectal neurons is homeostatically regulated across development and in response to short-term enhancement of visual stimulation, and this regulation is associated with changes in Na^+^ currents (Aizenman et al., 2003; Pratt and Aizenman, 2007). However, the molecular mechanisms by which Na^+^ currents are regulated to control the intrinsic excitability of tectal neurons, and whether regulation of K^+^ currents contribute to this control of excitability, are not fully understood. We therefore examined the relationship between intrinsic excitability of tectal neurons and the regulation of Na^+^ and K^+^ currents. To do this, we performed whole cell recordings on 65 deep-layer principal tectal neurons from tadpoles at developmental stage 49 using a whole brain *ex vivo* preparation (Figure 1A), as previously described (Wu et al., 1996). This developmental stage was chosen because the biophysical properties of tectal neurons are most heterogenous, with neurons mostly exhibiting low intrinsic excitability; yet, some highly excitable neurons are still observed (Ciarleglio et al., 2015).

**Figure 1:**
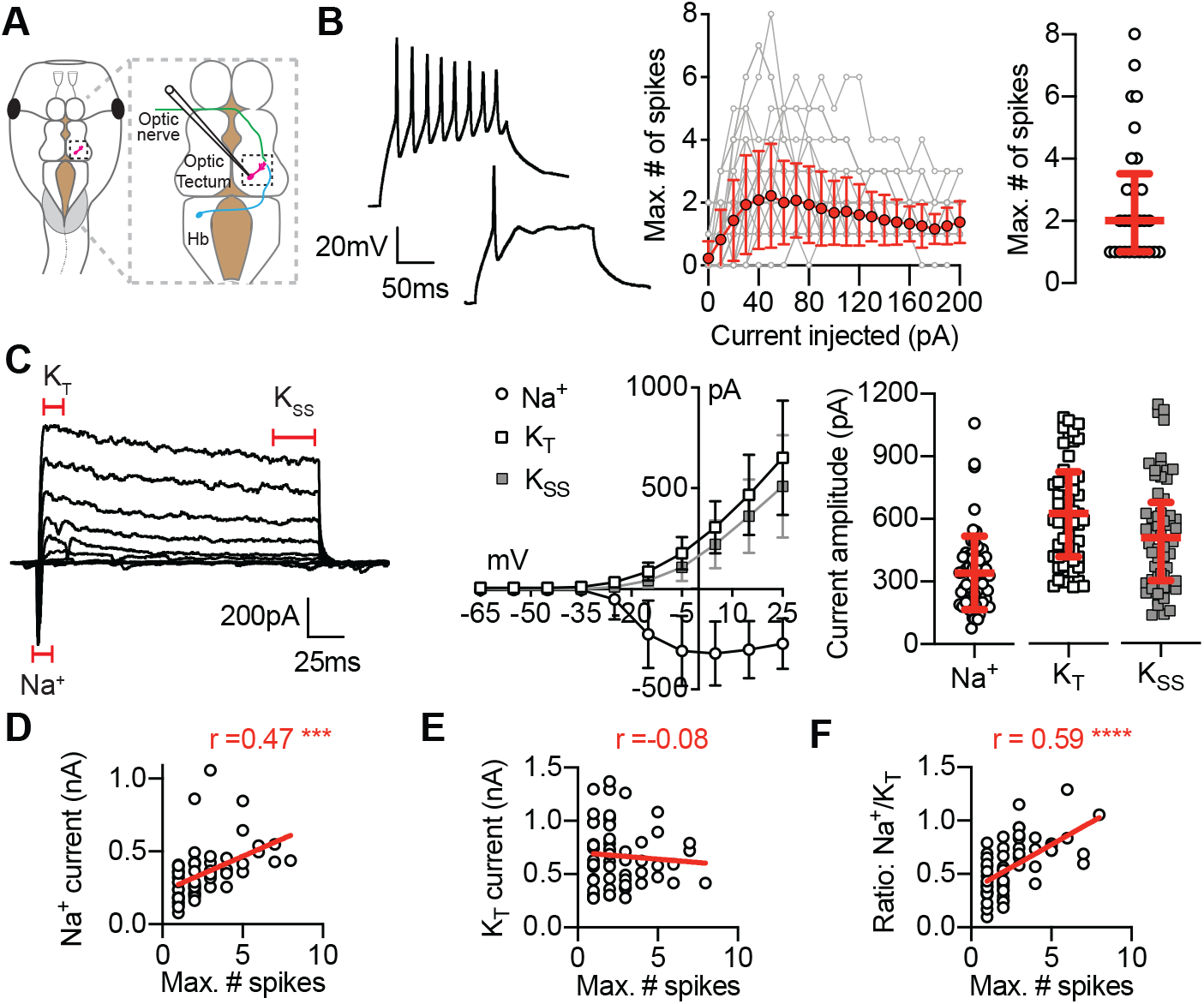
Intrinsic excitability of *Xenopus* tectal neurons is correlated with the amplitude of voltage-gated Na^+^ currents. **(A)** Diagram shows a *Xenopus* tadpole illustrating whole-cell recording from a neuron of the optic tectum that receives innervation from visual and mechanosensory inputs. **(B)** *Left*: Example current-clamp recordings showing the spiking response of two stage 49 tectal neurons held at −65mV to a 50pA current injection, illustrating the range of responses observed. *Middle*: Plot shows the number of spikes elicited in response to 0-200pA current injections for all cells analyzed (grey), and the average response (red, mean ± SD). *Right*: Maximum number of spikes (median and IQR, *n*=61) **(C)** *Left*: Example voltage-clamp recording from a tectal neuron held at −65mV in response to a series of depolarizing steps (−65mv to +25mV). Recording is leak subtracted to show only active currents. *Middle*: *I-V* plot shows average current amplitude for the Na^+^, the transient K^+^ and the steady state K^+^ currents. *Right*: Peak current amplitudes (mean ± SD; *n*=65). **(D-F)** Plots show Pearson correlations between increased intrinsic excitability (max. number of spikes) and the amplitude of the Na^+^ current, the transient K^+^ current, and the ratio of the Na^+^ current to transient K^+^ current. r values are Pearson correlation coefficients (\*\*\*\**p*<0.0001; \*\*\**p*=0.0002).

Intrinsic excitability was determined by measuring the ability of tectal neurons to generate action potentials in response to a series of 150ms depolarizing current steps ranging from 0-200pA (Figure 1B). As expected, we observed a broad range of values in the maximum number of spikes that tectal neurons could generate (1-8), with most neurons spiking once or twice (Figure 1B; median = 2 spikes, IQR 1-3 spikes, *n* = 61 cells). To examine the relationship between intrinsic excitability and Na^+^ and K^+^ currents in tectal neurons, we calculated the maximal current amplitudes for the voltage-gated Na^+^ current, the transient K^+^ current (K_T_), and the steady state K^+^ current (K_SS_) (Aizenman et al., 2003). Current amplitudes were measured by delivering a series of 150ms depolarizing steps starting from −65mv to +25mV to generate *I-V* plots, from which peak amplitudes were quantified (Figure 1C; Na^+^: 341.4 ± 176.1pA; K_T_: 666.9 ± 281.0pA; K_SS_: 517.1 ± 251.2pA; *n* = 65 cells). These intrinsic properties matched previous reports for stage 49 tectal neurons (Aizenman et al., 2002; Pratt and Aizenman, 2007; Ciarleglio et al., 2015), and showed a wide range of excitability, which allowed us to investigate the cellular basis for heterogeneity of intrinsic excitability within tectal neurons.

To determine the relationship between intrinsic excitability and other intrinsic properties of tectal neurons, we performed a multivariate analysis and calculated Pearson pairwise correlations between each of the biophysical properties measured for the 65 stage 49 tectal neurons (For complete correlation matrix see Figure 1 - figure supplement 1A). We found that intrinsic excitability, represented by the maximum number of spikes generated by each neuron, was most positively correlated with the peak amplitude of Na^+^ current (Figure 1D) and the Na^+^ to K_T_ current ratio (Figure 1F). In contrast, we observed no correlation between intrinsic excitability and the amplitude of the transient K^+^ current (Figure 1E). These data suggest that the intrinsic excitability of tectal neurons is primarily modulated by differences in Na^+^ currents and not K^+^ currents. Further supporting this idea, we found significant correlations between the amplitude of the Na^+^ current and characteristics of the first spike including threshold potential, rate of rise and spike width; however, we observed no correlation between characteristics of the first spike and the amplitude of either transient or steady state K^+^ currents (Figure 1 – figure supplement 1). These data support the hypothesis that the modulation of Na^+^ currents is the key mechanism for regulating the excitability of tectal neurons.

### Tectal neurons express distinct fast and persistent Na^+^ currents that are regulated with homeostatic changes in intrinsic excitability across development and in response to enhanced visual stimulation

How are voltage-gated Na^+^ currents regulated in order to modulate intrinsic excitability? Voltage-gated Na^+^ channels carry both fast and persistent Na^+^ currents, which have distinct roles in action potential generation and repetitive firing of neurons (French, 1990; Raman et al., 1997; Magistretti et al., 2006). The fast Na^+^ current mediates the upstroke of action potentials before becoming rapidly inactivated (Hodgkin and Huxley, 1952; Stuart and Sakmann, 1994). When Na^+^ channels fail to inactivate, even with prolonged depolarization, the resulting sustained current is termed a persistent Na^+^ current. This persistent Na^+^ current is activated at a subthreshold voltage range and can amplify a neurons response to synaptic input, enhancing its capacity to fire repetitively (Bant and Raman, 2010; Chen et al., 2011). We therefore hypothesized that tectal neurons may modulate fast and persistent Na^+^ currents to achieve homeostatic changes in intrinsic excitability. While fast Na^+^ currents are known to be modulated by activity (Aizenman et al., 2003; Pratt and Aizenman, 2007), the regulation of persistent Na^+^ currents has not been examined.

As the tectal circuitry matures between developmental stages 42 and 49, the intrinsic excitability of tectal neurons peaks at stage 46 preceding an overall increase in the level of excitatory synaptic input at this stage of circuit development (Figure 2A)(Aizenman et al., 2002; Ciarleglio et al., 2015). Therefore, to determine whether fast and persistent Na^+^ currents are regulated with changes in intrinsic excitability across development, we performed whole-cell recordings on tectal neurons from developmental stages 42, 46 and 49. Na^+^ currents were isolated in voltage-clamp recordings using a Tris-based internal saline solution to block outward K^+^ currents (Aizenman et al., 2003), which revealed distinct fast and persistent voltage-gated Na^+^ currents (Figure 2B). *I-V* plots were generated by measuring the peak amplitude of the fast and persistent Na^+^ currents in response to a series of 150ms depolarizing steps ranging from −65 to 25mV (Figure 2C-D). Interestingly, increasing the time of the depolarizing step from 150 to 400ms had no effect on the amplitude of the persistent Na^+^ current (*data not shown*), which is consistent with persistent Na^+^ currents having ultra-slow inactivation kinetics in tectal neurons. When we compared peak amplitudes of the fast and persistent voltage-gated Na^+^ currents across development stages, we observed a transient peak at stage 46 (Figure 2E-F and Figure 2 - figure supplement 2), when intrinsic excitability of tectal neurons is highest (Pratt and Aizenman, 2007). These data confirm that fast Na^+^ currents are regulated with changes in intrinsic excitability of tectal neurons during development, and extend these findings to show that a persistent Na^+^ current is similarly transiently upregulated at stage 46, suggesting that fast and persistent Na^+^ currents are co-regulated with changes in intrinsic excitability.

**Figure 2:**
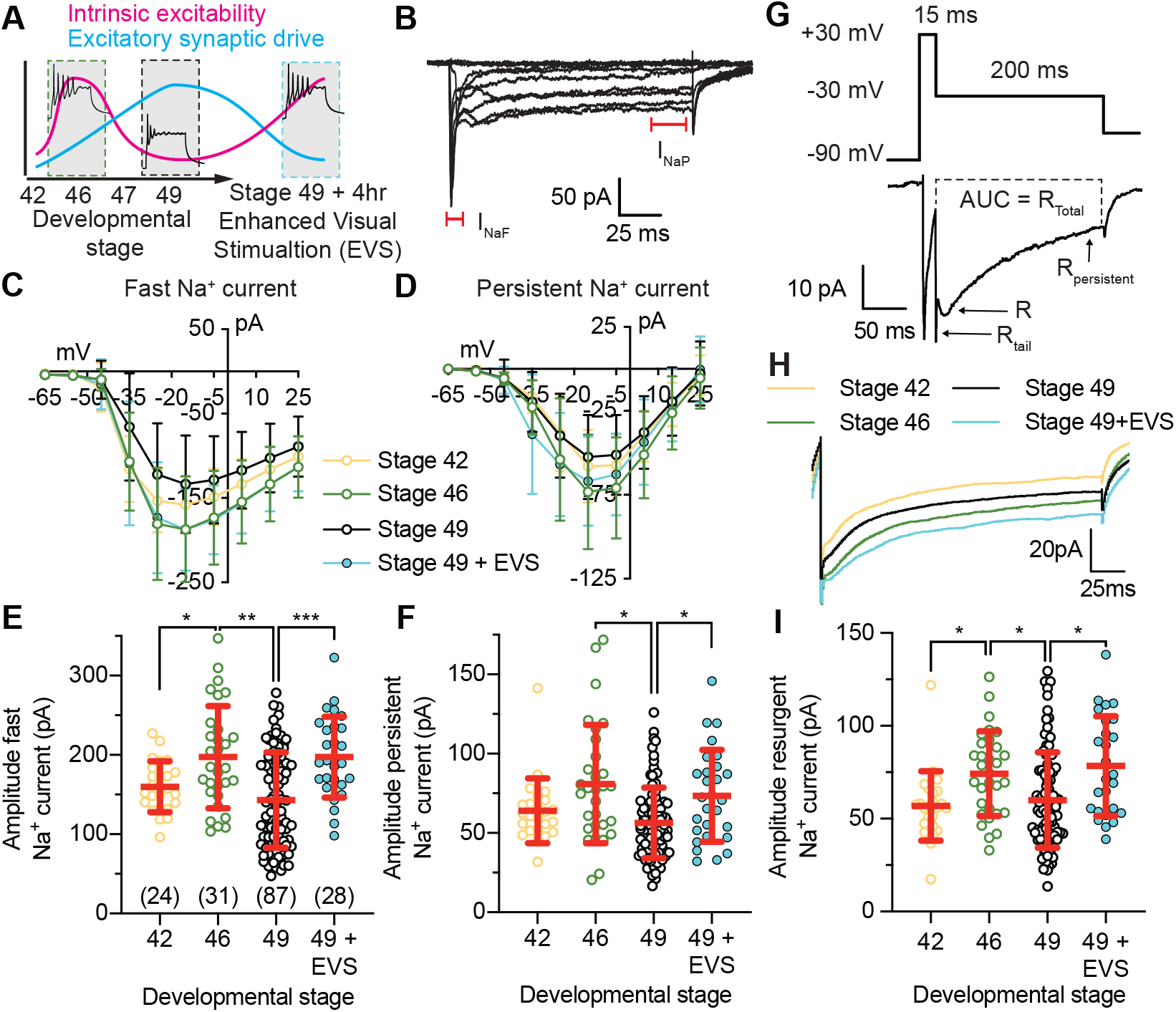
Tectal neurons express fast and persistent voltage-gated Na^+^ currents that are regulated with changes in intrinsic excitability. **(A)** Schematic depicts how tectal neurons homeostatically adapt intrinsic excitability in response to changing excitatory synaptic drive across development or in response to 4hr exposure to enhanced visual stimulation to maintain a broad dynamic range and, thereby, conserve input-output function as the tectal circuitry changes. **(B)** Example voltage-clamp recording from a tectal neuron held at −65mV in response to a series of 150ms depolarizing steps (−65mV to +25mV) using a Tris-based internal saline solution to block outward K^+^ currents. Recording is leak subtracted to show only active currents. Fast (I_NaF_) and persistent (I_NaP_) Na^+^ currents are indicated on the recording. **(C-D)** Averaged *I-V* plots from tectal neurons sampled across development, and at stage 49 in response to 4hrs of EVS to increase intrinsic excitability, for the fast and persistent Na^+^ currents. **(E-F)** Quantifications of peak amplitudes for the fast and persistent Na^+^ currents (median and IQR. **(G)** Example resurgent current recording from a tectal neuron held at −65mV that was hyperpolarized to −90mV before stepping to +30mV for 15ms to open voltage-gated Na^+^ channels. Resurgent Na^+^ currents were then recorded by stepping to −30mV for 200ms, with distinct tail resurgent (R_Tail_), resurgent (R) and persistent resurgent (R_Persistent_) currents identified. Recordings were obtained using a Tris-based internal saline solution to block outward K^+^ currents, and leak subtracted to show only active currents. **(H)** Averaged resurgent current traces obtained from tectal neurons between developmental stages 42-49, and at stage 49 following exposure to 4hrs of enhanced visual stimulation (EVS). **(I)** Peak resurgent current amplitude across development and in response to 4hrs of EVS. Groups were compared using a Welch’s ANOVA test with Dunnett T3 test for multiple comparisons. *n* values are shown in E.

The finding that fast and persistent Na^+^ currents are transiently increased at stage 46, when intrinsic excitability is highest, supported the hypothesis that tectal neurons modulate fast and persistent Na^+^ currents together to regulate intrinsic excitability. It is not clear, however, whether this transient increase in fast and persistent Na^+^ currents observed at stage 46 is a function of tectal neuron development, or if this represents a mechanism for the homeostatic regulation of intrinsic excitability in tectal neurons. If the regulation of voltage-gated fast and persistent Na^+^ currents is a mechanism to homeostatically control intrinsic excitability, then it would be predicted that established protocols for inducing homeostatic increase in intrinsic excitability would also cause an increase in fast and persistent Na^+^ currents. To test this prediction, we exposed stage 49 tadpoles to 4hrs of enhanced visual stimulation (EVS), which triggers increased intrinsic excitability of tectal neurons as a homeostatic response to decreased excitatory synaptic drive (Figure 2A). To expose tadpoles to EVS, freely swimming stage 49 tadpoles were placed for 4hrs in a custom-built chamber containing a LED light array that flashed at 1Hz in sequence to simulate a motion stimulus (Sin et al., 2002; Ciarleglio et al., 2015). When we compared the amplitude of the fast and persistent voltage-gated Na^+^ currents between tectal neurons from naïve stage 49 tadpoles or stage 49 tadpoles exposed to 4hrs of EVS, we observed a significant increase in the amplitude of both the fast and persistent Na^+^ currents (Figure 2C-F and Figure 2 - figure supplement 2). These data show that tectal neurons can modulate fast and persistent Na^+^ currents with homeostatic changes in intrinsic excitability, providing further evidence that the regulation of Na^+^ currents underlies the homeostatic regulation of intrinsic excitability in tectal neurons.

### Tectal neurons express resurgent Na^+^ currents that are regulated with homeostatic changes in intrinsic excitability across development and in response to enhanced visual stimulation

In addition to fast and persistent Na^+^ currents, voltage-gated Na^+^ channels can also mediate a resurgent Na^+^ current that is known to facilitate repetitive firing in many types of neurons (Raman et al., 1997; Cummins et al., 2005; Bant and Raman, 2010; Theile and Cummins, 2011; Browne et al., 2017). Voltage-gated Na^+^ channels opened by membrane depolarization rapidly inactivate to prevent excessive Na^+^ influx. However, in cells that express the resurgent Na^+^ current, a fraction of Na^+^ channels do not inactivate via the channel’s own inactivation domain but are arrested in an open state by open-channel block, typically by an accessory subunit, which prevents the fast inactivation gate from binding and inactivating open channels. When this occurs, repolarization of the neuron causes the open channel block to be expelled, allowing the resurgent Na^+^ current to flow until the channel is closed. We tested whether tectal neurons express a resurgent Na^+^ current and whether this resurgent Na^+^ current could contribute to the homeostatic control of intrinsic excitability.

To measure resurgent Na^+^ currents, we first performed a test pulse to 0mV to measure the fast Na^+^ current and then hyperpolarized the neuron to −90mV for 500ms to promote a shift in inactivated Na^+^ channels to a closed state. We then depolarized cells to 30mV for 15ms to open Na^+^ channels and induce open-channel block, before hyperpolarizing to −30mV to generate resurgent Na^+^ currents (Figure 2G). We identified distinct components of the resurgent current (see labelled example trace in Figure 2G). We observed a tail resurgent current (R_Tail_) that occurred ∼1ms after cells were repolarized to −30mV (Rise time [0-100%] = 0.92ms ± 0.39ms; *n* = 156 cells across all developmental stages). We termed this current a tail resurgent Na^+^ current as the amplitude of the current showed an almost linear voltage-dependency with the voltage of the resurgent step (Figure 2 - figure supplement 1D-F), and decreased as the time of the depolarizing step increased (Figure 2 - figure supplement 1G-I), consistent with characteristics of tail currents. We also observed a resurgent Na^+^ current (R) that occurred following the tail current at ∼7.5ms (Rise time [0-100%] = 7.50ms ± 9.64ms) and slowly decayed to a persistent state with a half-life of ∼24ms (Decay time [100-50%] = 23.99ms ± 13.64ms). This current most closely resembled resurgent currents initially described in rodent Purkinje neurons (Raman et al., 1997; Raman and Bean, 2001), and exhibited voltage dependency that peaked around −60mV as the voltage of the resurgent step was varied from +20mV to −90mV (Figure 2 - figure supplement 1D-F), decreased as the time of the depolarizing step increased (Figure 2 - figure supplement 1G-I), and increased as the voltage of the depolarizing step was increased from −30 to +30mV (Figure 2 - figure supplement 1J-L), which is consistent with descriptions of resurgent Na^+^ currents. We also observed a persistent Na^+^ resurgent current (R_Persistent_; average current over the final 10ms of the −20mV step). This current was highly voltage-dependent, as illustrated by a peak conductance between 0 and −30mV (Figure 2 - figure supplement 1D-F); however, this current was not affected by changes in the time or voltage of the depolarizing step (Figure 2 - figure supplement 1G-L). Similar to what was observed for the persistent Na^+^ current, we observed no effect on the persistent resurgent current of increasing the time of the resurgent step to 400ms (data not shown), which may suggest the presence of Na^+^ channels with ultra-slow inactivation kinetics. We also measured the total resurgent Na^+^ current, by calculating the total charge (pA*s) over the entire 200ms step. Similar to the persistent resurgent current, the total resurgent current measurement showed dramatic voltage-dependency in response to varying the voltage of the resurgent step, also peaking between 0 and −30mV (Figure 2 - figure supplement 1D-F). However, there was little effect of altering the time or voltage of the depolarizing step (Figure 2 - figure supplement 1G-L). These data are the first description of a resurgent Na^+^ current in *Xenopus* tectal neurons, providing another mechanism by which Na^+^ currents could be regulated to control intrinsic excitability.

If the presence and regulation of a resurgent Na^+^ current is a mechanism used by tectal neurons to regulate homeostatic control of intrinsic excitability, then we would predict that the resurgent current would be regulated with homeostatic changes in intrinsically excitability across development and in response to EVS. Consistent with changes we observed in fast and persistent Na^+^ currents, when resurgent currents were compared across development we found that the resurgent Na^+^ current peaked at stage 46 and was increased in response to 4hrs of EVS exposure (Figure 2I and Figure 2 - figure supplement 2). Furthermore, we observed that exposure to EVS caused an increase in the total resurgent current (Figure 2 - figure supplement 1C, and figure supplement 2), but had no effect on the tail or persistent resurgent Na^+^ currents (Figure 2 - figure supplement 1A-B and figure supplement 2), supporting the view that changes in the resurgent Na^+^ current correlate well with changes in intrinsic excitability. Taken together, our data suggest that tectal neurons regulate fast, persistent and resurgent Na^+^ currents together to control homeostatic changes in intrinsic excitability.

### Persistent and resurgent Na^+^ currents, but not the fast Na^+^ current, are insensitive to TTX

*Xenopus* tectal neurons express an array of voltage-gated currents including a TTX-sensitive fast Na^+^ current, a persistent Na^+^ current, transient and steady state K^+^ currents, and small Ca^2+^ current (Aizenman et al., 2003; Pratt and Aizenman, 2007; Hamodi and Pratt, 2014). Our findings had shown that fast, persistent and resurgent Na^+^ currents are correlated with changes in intrinsic excitability, suggesting that the regulation of Na^+^ currents may underlie the capacity of tectal neurons to homeostatically control their intrinsic excitability. Nothing is known, however, about the function, molecular characterization and pharmacology of these persistent and resurgent Na^+^ current in these cells. We therefore performed a series of experiments utilizing either ion substitution or specific channel inhibitors to characterize the ionic permeability and channel types that contribute to the persistent and resurgent Na^+^ currents observed in tectal neurons.

To confirm that persistent and resurgent Na^+^ currents identified in this study were the result of Na^+^ influx, we performed an ion substitution experiment and replaced extracellular Na^+^ with N-Methyl-D-glucamine (NMDG), an impermeant organic monovalent cation that abolishes inward Na^+^ currents (Blair and Bean, 2002). When we compared fast, persistent and resurgent Na^+^ currents in control stage 49 tectal neurons (black traces in Figure 3A,B,E) with stage 49 tectal neurons in NMDG external (grey traces in Figure 3A,B,E), we observed that extracellular Na^+^ is necessary for fast, persistent and resurgent Na^+^ currents, with substitution of extracellular Na^+^ with NMDG abolishing these Na^+^ currents (Figure 3A-F and Figure 3 - figure supplement 2A-C). However, a small presumptive Ca^2+^ current remains (Hamodi and Pratt, 2014). Importantly, replacing extracellular Na^+^ with NMDG had no effect on voltage-gated K^+^ currents (Figure 3 - figure supplement 2A-C), illustrating the specificity of the effect of Na^+^ substitution on voltage-gated Na^+^ currents. These data provide further evidence that voltage-gated sodium channels mediate the fast, persistent and resurgent Na^+^ currents observed in *Xenopus* tectal neurons.

**Figure 3:**
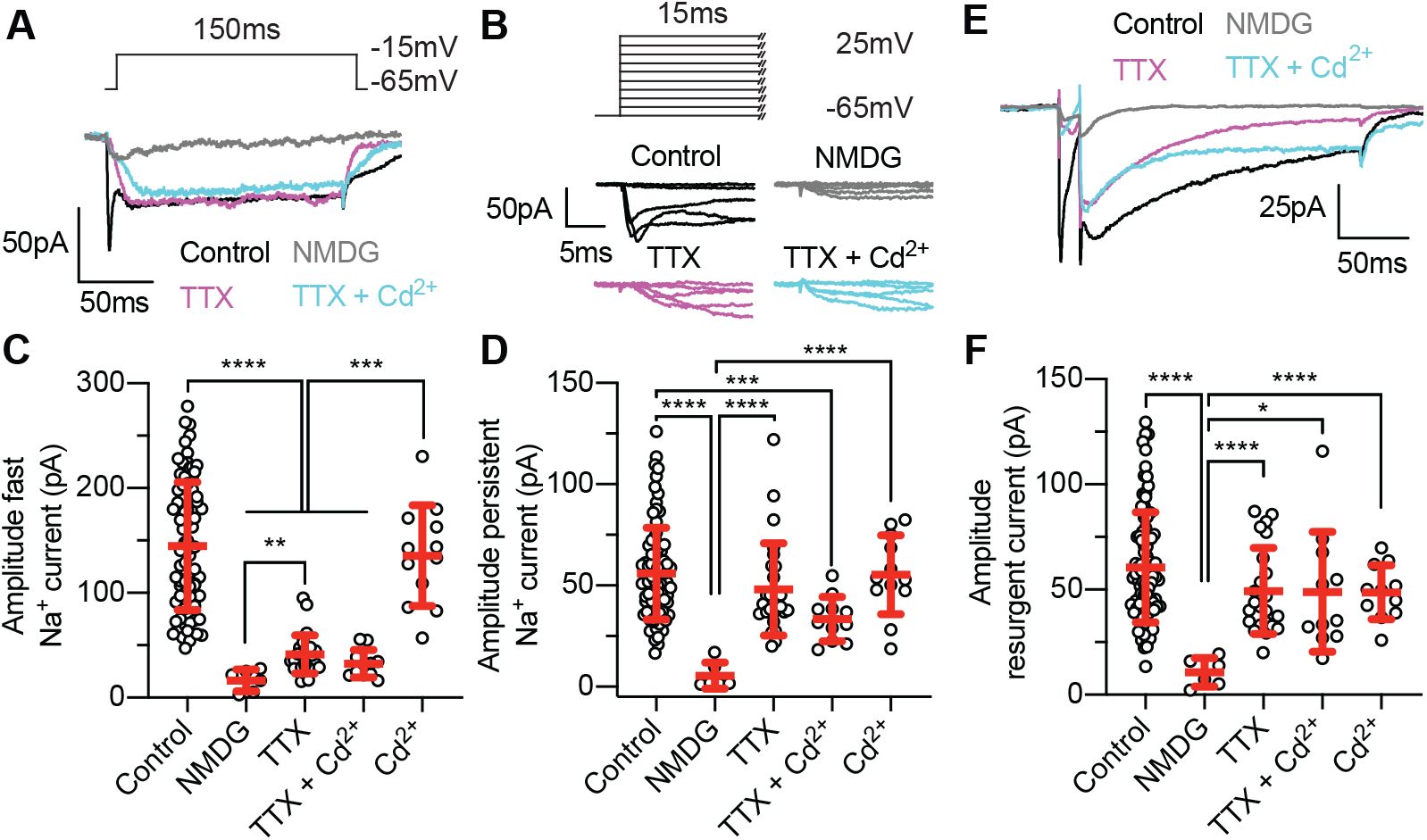
The persistent and resurgent Na^+^ currents, but not the fast Na^+^ current, are insensitive to TTX. **(A)** Example recordings from tectal neurons showing a single depolarizing step to −15mV to activate voltage-gated Na^+^ currents. Recordings were obtained from a stage 49 tectal neurons in control conditions (black), in zero Na^+^ external solution (NMDG) to block all inward Na^+^ currents (grey), in 1µM TTX to block TTX-sensitive Na^+^ currents (magenta), or in 100nM Cd^2+^ to block Ca^2+^ currents (cyan). Note that TTX attenuates fast but not persistent Na^+^ currents, whereas blocking all Na^+^ influx by replacing external Na^+^ with NMDG attenuates both fast and persistent currents to reveal a small presumptive Ca^2+^ current. **(B)** Magnification of the initial 15ms of the response of cells shown in (A) to a series of voltage steps (−65mV to +25mV) highlighting the effect of each treatment on the fast Na^+^ current. **(C-D)** Quantification of peak amplitudes for the **(C)** fast and **(D)** persistent Na^+^ currents. **(E)** Example resurgent current traces obtained from stages 49 tectal neurons. **(F)** Quantification of peak amplitudes for the resurgent Na^+^ currents. *n* values for C,D & F were 87 (stage 49 control), 15 (NMDG), 27 TTX, 11 (TTX + Cd^2+^), and 12 (Cd^2+^).

Previous studies have shown that the fast Na^+^ current in *Xenopus* tectal neurons is sensitive to the Na^+^ channel blocker tetrodotoxin (TTX) (Aizenman et al., 2003). We therefore predicted that the persistent and resurgent Na^+^ currents would also be sensitive to TTX. To test this prediction we measured the fast, persistent and resurgent Na^+^ currents in the presence of 1µM TTX (magenta traces in Figure 3A,B,E). Surprisingly, while TTX attenuated the fast Na^+^ current as expected, there was no effect of TTX on the amplitude of the persistent or resurgent Na^+^ currents (Figure 3A-F and Figure 3 - figure supplement 1). Increasing the concentration of TTX to 30µM had no effect on the persistent or resurgent Na^+^ currents (Figure 3 - figure supplement 2D-F), further suggesting that persistent and resurgent Na^+^ currents are TTX-insensitive in tectal neurons. Furthermore, when control or TTX-exposed tectal neurons were exposed to a depolarizing voltage ramp from −90mV to 30mV (0.8mV/s), we observed distinct voltage-dependencies for the TTX-sensitive fast Na^+^ current and the TTX-insensitive persistent Na^+^ current (Figure 3 - figure supplement 2G), which suggests that persistent Na^+^ currents may be carried by a distinct subset of TTX-insensitive voltage gated Na^+^ channels in tectal neurons.

We had observed the presence of TTX-insensitive persistent and resurgent Na^+^ currents. In other systems, persistent and resurgent Na^+^ currents have been effectively targeted using the Na^+^ channel modulators Riluzole and GS967 (Urbani and Belluzzi, 2000; Baker et al., 2018). We therefore asked whether these Na^+^ channel modulators could affect Na^+^ currents in *Xenopus* tectal neurons. However, we observed no effect on fast, persistent or resurgent Na^+^ currents of these treatments (Figure 3 - figure supplement 2D-F), suggesting that these Na^+^ channel modulators have no effect on Na^+^ currents in *Xenopus* tectal neurons.

When Na^+^ currents were abolished by substituting extracellular Na^+^ for NMDG we observed a small, presumptive Ca^2+^ current (Figure 3A-B). We therefore asked whether the TTX-insensitive component of persistent and resurgent Na^+^ currents represents Ca^2+^ influx via voltage-gated Ca^2+^ channels. To test this, we blocked voltage-gated Ca^2+^ channels with 100nM Cd^2+^ in the presence or absence of TTX (cyan traces in in Figure 3A,B,E). Crucially, we observed no effect of Cd^2+^ effect on the amplitude of the fast, persistent or resurgent Na^+^ current (Figure 3A-F and Figure 3 - figure supplement 1). Other pharmacological blockers of Ca^2+^ channels similarly did not affect the Na^+^ currents (Figure 3 - figure supplement 2D-F). These findings are consistent with the TTX-insensitive component of the persistent and resurgent Na^+^ currents being mediated by Na^+^ influx via voltage-gated Na^+^ channels.

We had observed persistent and resurgent Na^+^ currents that were insensitive to TTX and that remained following blockage of voltage-gated Ca^2+^ channels. Therefore, we also tested whether this influx could be the result of inward flux of K^+^ ions. To do this, we examined the effect of TEA external on the fast, persistent, and resurgent Na^+^ currents. Unsurprisingly, we observed no effect on fast, persistent or resurgent Na^+^ currents TEA external (Figure 3 - figure supplement 2D-F), which provides further evidence that the described TTX-insensitive persistent and resurgent Na^+^ currents are indeed Na^+^ currents. Taken together, these experiments confirm that tectal neurons express a TTX-sensitive fast Na^+^ current and extend this finding to demonstrate the presence of TTX-insensitive component of the persistent and resurgent Na^+^ currents.

### Inhibition of Na^+^ channel subtype Na_v_1.6 attenuates resurgent Na^+^ currents and decreases the intrinsic excitability of tectal neurons

We had identified that *Xenopus* tectal neurons express TTX-sensitive fast Na^+^ current and TTX-insensitive persistent and resurgent Na^+^ currents. The molecular characterization of the persistent and resurgent Na^+^ currents, however, remained unclear. Na^+^ channels are comprised of alpha and beta subunits, the alternate expression of which confers Na^+^ channels with distinct cellular expression profiles, subcellular localization, conduction properties, and responses to Na^+^ channel activators and inhibitors (Savio-Galimberti et al., 2012). Alpha subunits (Na_v_1.1-Na_v_1.9) are pore-forming components of Na^+^ channels, whereas beta subunits (Na_v_1β-Na_v_4β) are accessory subunits that non-covalently bind and regulate the expression, cellular localization and gating properties of Na^+^ channels (O’Malley and Isom, 2015). A search of the *Xenopus laevis* genome (Xenbase: *X. laevis* version 9.2 on JBrowse) revealed that *Xenopus* express genes encoding for the neuronal channel subtypes Na_v_1.1, Na_v_1.2 and Na_v_1.6; as well as the accessory beta subunit Na_v_4β, which has a well described role in regulating resurgent Na^+^ currents (Bant and Raman, 2010). Although Na_v_1.6 is TTX-sensitive in mammalian neurons (Lee and Ruben, 2014), it is evolutionarily distinct from Na_v_1.1 and Na_v_1.2 (Zakon, 2012), and has been shown to mediate persistent and resurgent Na^+^ currents (Raman et al., 1997; Bant and Raman, 2010; Patel et al., 2015), making it a candidate to mediate TTX-insensitive Na^+^ currents in tectal neurons. We therefore hypothesized that Na_v_1.6 could be the molecular determinate of persistent and resurgent Na^+^ currents in *Xenopus* tectal neurons.

A recent study described a novel, specific Na_v_1.6 inhibitor, MV1312, which shows a 5-6 fold sensitivity for Na_v_1.6 over Nav1.1-1.7, and comparable blocking efficiency for Na_v_1.8 (Weuring et al., 2020). MV1312 has been shown to rescue seizure behavior in a zebrafish model of Dravet syndrome where a loss of Na_v_1.1 function results inappropriate compensation by Na_v_1.6 and epileptogenesis (Weuring et al., 2020). We therefore asked whether MV1312 can block Na^+^ currents in *Xenopus* tectal neurons. To measure the effect of MV1312 on fast, persistent and resurgent Na^+^ currents, we performed whole cell recordings with a Tris-based internal solution and washed in 5µM MV1312. Interestingly, when we acutely washed in MV1312, we observed no effect of MV1312 on fast, persistent or resurgent Na^+^ currents (Figure 4 - figure supplement 1 and 2). However, when current recordings were preceded by a 5s depolarizing step to 0mV to open sodium channels, we found that MV1312 caused a significant attenuation of all Na^+^ currents, with the largest effect observed for the resurgent Na^+^ current (Figure 4A-D and Figure 4 - figure supplement 2). These findings suggest that MV1312 blocks Na^+^ channels in the open configuration with slow kinetics, and that Na_v_1.6 is a major contributor to fast, persistent and resurgent Na^+^ currents in tectal neurons.

**Figure 4:**
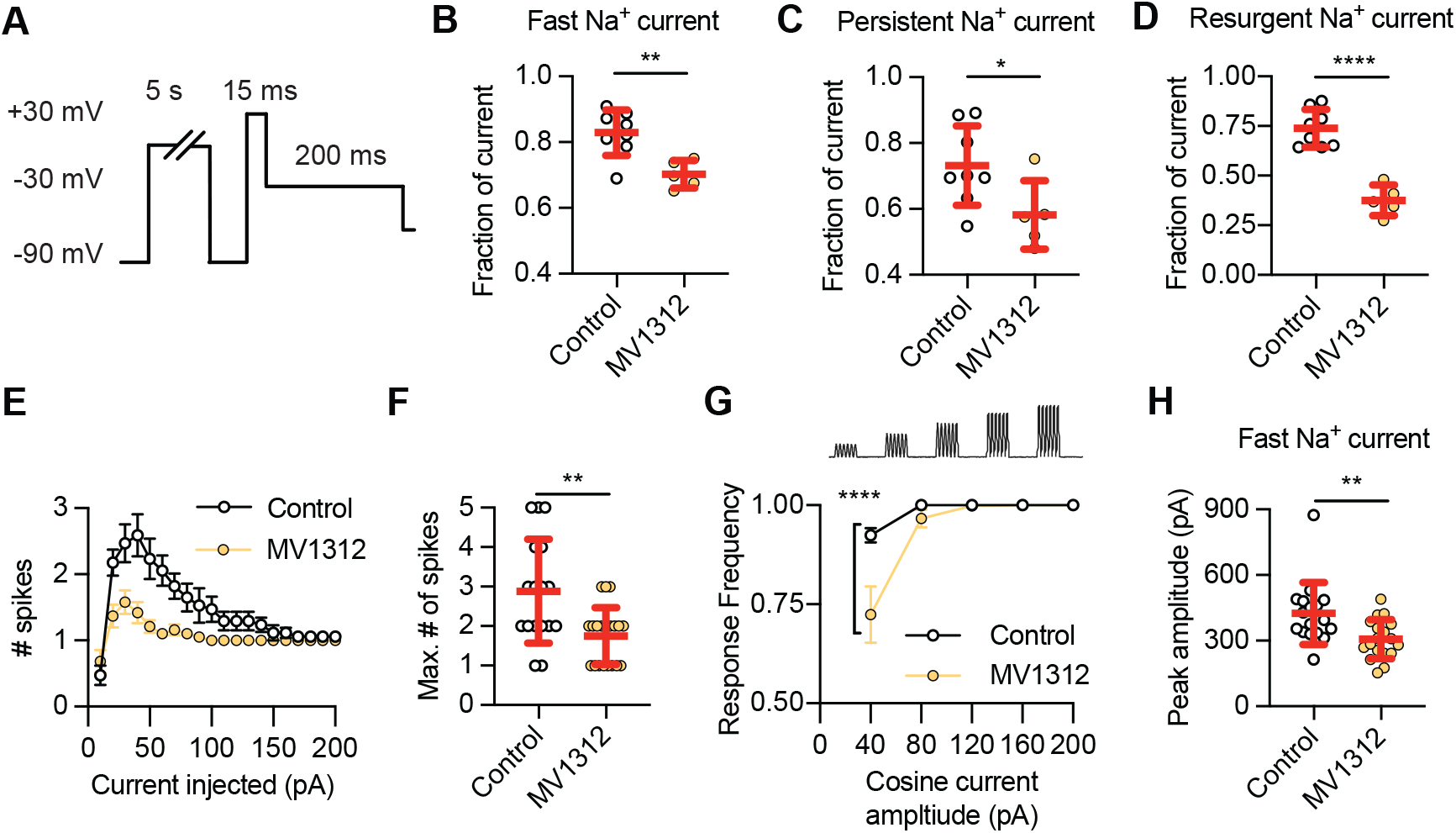
The Na_v_1.6 specific inhibitor MV1312 decreases intrinsic excitability by reducing fast, persistent and resurgent Na^+^ currents in a use-dependent manner. **(A-D)** Acute wash in of the specific Na_v_1.6 inhibitor MV1312 reduces Na^+^ current amplitude in a use-dependent manner. **(A)** When acute wash in of MV1312 is combined with a 5s depolarizing step to 0mV preceding recording of the resurgent Na^+^ currents, there is a reduction in the **(B)** fast Na^+^ current, **(C)** persistent Na^+^ current, and **(D)** the resurgent Na^+^ current. **(E-F)** MV1312 decreases intrinsic excitability of tectal neurons by attenuating Na^+^ current amplitude. The effect of 5µM MV1312 was observed by measuring **(E)** spikes generated by current injection, **(F)** the maximum number of spikes generated by current injection, **(G)** the capacity of cells to spike in response to a cosine injection of current (40 to 200pA), and **(H)** by measuring peak amplitude of the fast Na^+^ current.

If Na_v_1.6 mediated Na^+^ currents are important regulators of intrinsic excitability in tectal neurons, then it would be predicted that acute inhibition of Na_v_1.6 channels would reduce intrinsic excitability. We therefore tested the effect of MV1312 wash in on intrinsic excitability of tectal neurons by performing whole cell recordings with a K^+^-based internal solution. As tectal neurons at stage 49 inherently exhibit lower intrinsic excitability, we performed recordings at stage 47/48 when the intrinsic excitability is higher (Pratt et al., 2008; Ciarleglio et al., 2015). As expected, the intrinsic excitability of control tectal neurons was increased in this population of cells (Figure 4E-F; median = 3 spikes, IQR 2-4 spikes, *n* = 17 cells) compared to neurons at stage 49 (compare with Figure 1B). Significantly, when we compared the intrinsic excitability of control tectal neurons with tectal neurons exposed to MV1312, we observed a decrease in intrinsic excitability (Figure 4E-F; median = 2 spikes, IQR 1-2 spikes, *n* = 20 cells; *p*=0.0047). These data suggest that Na^+^ currents carried by Na_v_1.6 channels are important for repetitive firing in tectal neurons.

To further study how Na_v_1.6 channel inhibition affects the capacity of tectal neurons to fire repetitively, we tested whether MV1312 effects the capacity of tectal neurons to generate action potentials in response to a cosine-shaped injection of current with different amplitudes. For these experiments, cells were exposed to repeated 200ms cosine current injections at 30Hz, with an increasing current amplitude from 40 to 200pA (Figure 4G; top). The response frequency for each cosine current amplitude was calculated by measuring whether a spike was generated by each individual cosine wave. Significantly, while control cells were able to faithfully generate spikes in response to cosine current injections at all current amplitudes, MV1312-exposed tectal neurons had a lower response frequency at 40pA (Figure 4G), consistent with MV1312 preventing repetitive firing by blocking Na_v_1.6-mediated Na^+^ currents. Furthermore, we observed a decrease in amplitude of the fast Na^+^ current in MV1312-exposed tectal neurons compared with control neurons (Figure 4H). Taken together, these data suggest that Na_v_1.6 channels contribute to fast, persistent, and resurgent Na^+^ currents and that these currents are important regulators of intrinsic excitability.

### Expression of Na^+^ channel subtypes are differentially regulated in the *Xenopus* optic tectum during key developmental time windows and in response to changes in network activity

Given that changes in Na^+^ current amplitudes and kinetics can be attributed to a change in the activity of distinct Na^+^ channel subtypes, and that we had observed that inhibition of Na_v_1.6 channels attenuates Na^+^ currents and decreases intrinsic excitability of tectal neurons, we next asked whether the expression of Na^+^ channel genes is regulated with homeostatic changes in Na^+^ current amplitudes and intrinsic excitability across developmental stages examined in this study, or in response to exposure to short-term patterned sensory experience.

To determine whether the expression of Na^+^ channel genes correlates with changes in Na^+^ current amplitudes and intrinsic excitability, we used real-time qPCR to quantify the expression of Na^+^ channel genes Na_v_1.1, Na_v_1.2, Na_v_1.6 and Na_v_4β in the optic tectum across development (Figure 5B). We found that expression levels of Na_v_1.1 and Na_v_1.6 in the optic tectum were regulated with development, but that the expression levels of Na_v_1.2 and Na_v_4β remained unchanged (Figure 5B). We observed a ∼2-fold increase in the level of Na_v_1.6 expression between stage 42 and stage 46 when Na^+^ currents and intrinsic excitability peaks (st 42: 1.00 ± 0.09, st 46: 1.71 ± 0.22; *p*=0.0350), with the level of expression at stage 49 being not significantly different from stages 46 (st 49: 1.40 ± 0.32; *p*=0.1932) or stages 42 (*p*=0.1932). In contrast, while the expression of Na_v_1.1 increased with developmental stage, as observed as a ∼2-fold increase in expression between stage 42 and stage 49 (st 42: 1.00 ± 0.12, st 49: 1.84 ± 0.31; *p*=0.0151), there was no significant change in the level of Na_v_1.1 expression between stages 42 and stage 46 (st 46: 1.57 ± 0.25; *p*=0.054), or between stage 46 and stage 49 (*p*=0.211). These data suggest that Na_v_1.6 expression correlates with changes in Na^+^ currents and intrinsic excitability across development, implying an important role for Na_v_1.6 in regulating developmental changes in Na^+^ current amplitude and intrinsic excitability as tectal neurons mature. In contrast, while Na_v_1.1 upregulation occurs with the maturation of the tectal circuitry, it was not directly correlated with changes in intrinsic excitability.

**Figure 5:**
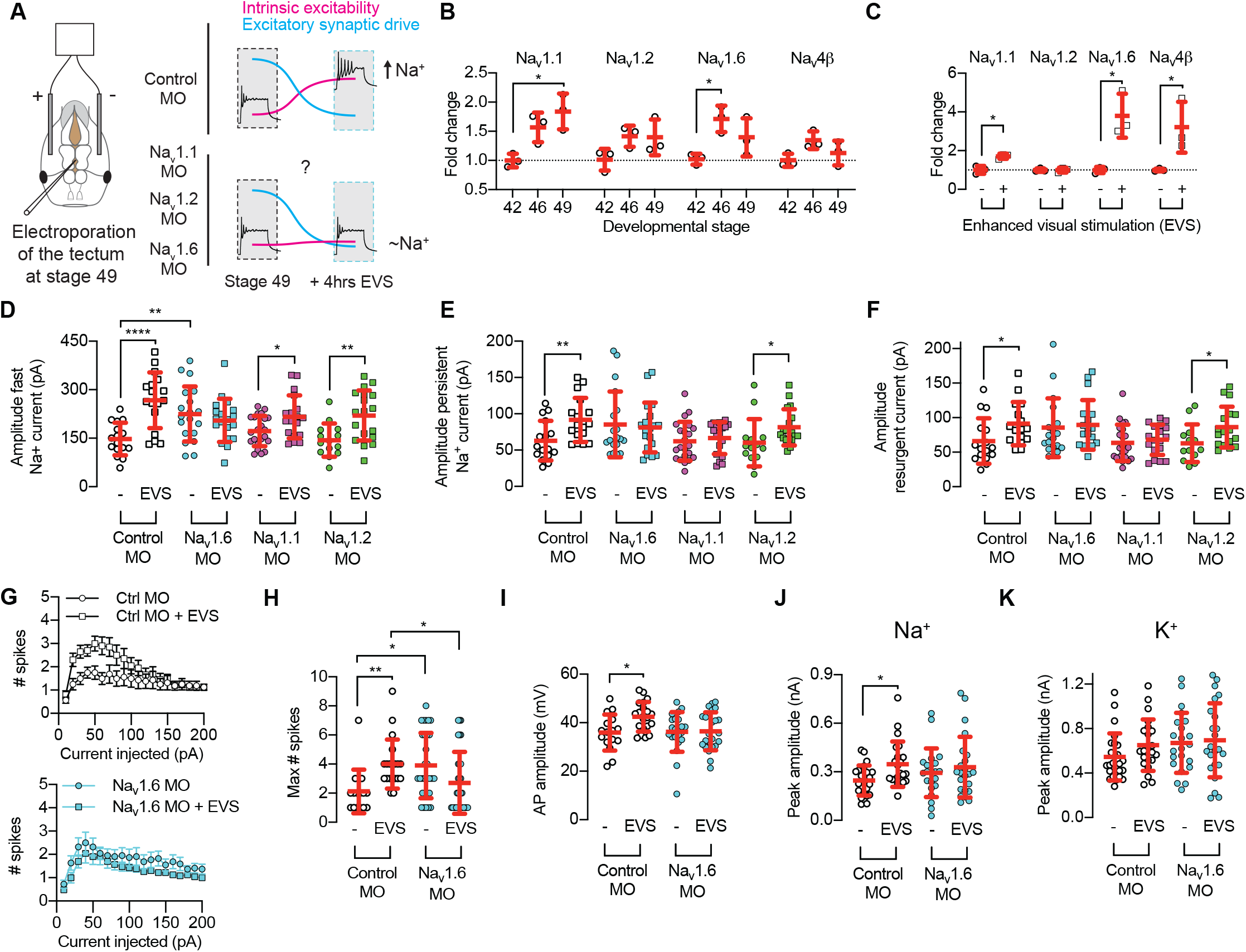
Knockdown of Na_v_1.6 attenuates network activity-dependent homeostatic increase in Na^+^ currents. **(A)** Schematic illustrates bulk electroporation of stage 49 tadpoles with morpholino oligonucleotides (MO), including a control MO and MOs targeting sodium channel alpha-subunit genes Na_v_1.1, 1.2 and 1.6. In normal conditions, exposure of stage 49 tadpoles to 4hrs of enhanced visual stimulation (EVS) decreases excitatory synaptic drive and triggers a compensatory increase in intrinsic excitability via an increase in sodium currents. **(B)** Expression of selected Na^+^ channel alpha and beta subunits in the optic tectum across development stages 42, 46 and 49. The expression of Na^+^ channel alpha and beta subunits genes was normalized to a housekeeper (RSP13), before determining the fold change in expression from stage 42. **(C)** Expression of selected Na^+^ channel alpha and beta subunits in the optic tectum at stage 49 in response to 4hr exposure to enhanced visual stimulation (EVS). Expression is shown as a fold change in normalized Na^+^ channel alpha and beta subunits gene expression compared to naïve stage 49 control tadpoles. **(D-F)** Exposure of stage 49 tadpoles to 4hrs of EVS triggers an increase in the amplitude of fast persistent and resurgent Na^+^ currents, which is attenuated by the knockdown of Na_v_1.6. **(G-K)** Exposure of stage 49 tadpoles to 4hrs of EVS triggers an increase in the amplitude of fast persistent and resurgent Na^+^ currents, which is attenuated by the knockdown of Na_v_1.6.

To test whether the upregulation of Na_v_1.6 expression is also a mechanism for homeostatically increasing Na^+^ current amplitude and intrinsic excitability, we tested whether Na_v_1.6 expression was also upregulated in the optic tectum of tadpoles exposed to EVS (Figure 5C). We found that the level of expression of Na_v_1.6 was increased ∼4-fold in the optic tectum of tadpoles exposed to 4hrs of EVS compared to naïve stage 49 controls (Ctrl: 1.00 ± 0.18, EVS: 3.80 ± 1.13; *p*=0.0135). We also observed a large ∼3-fold increase in the expression of Na_v_4β (Ctrl: 1.00 ± 0.07, EVS: 3.21 ± 1.31; *p*=0.0434). Taken together, these data provide further evidence that tectal neurons homeostatically control their intrinsic excitability by regulating the amplitude of Na^+^ currents through changes in gene expression levels of Na_v_1.6 channel subunits. We also observed a small but significant increase in the level of Na_v_1.1 expression in the optic tectum following EVS exposure (Ctrl: 1.00 ± 0.23, EVS: 1.71 ± 0.13; *p*=0.0108), suggesting that expression of Na_v_1.1 may be also increased to modulate the overall level of Na^+^ currents during the homeostatic increase in intrinsic excitability.

### Regulation of Na^+^ channel subtypes Na_v_1.1 and Na_v_1.6 mediate homeostatic changes in Na^+^ currents to control the intrinsic excitability of tectal neurons

We had observed changes in the expression of Na_v_1.1 and Na_v_1.6 channels across development and in response to EVS, suggested a role for these Na^+^ channel subtypes in the homeostatic regulation of intrinsic excitability. We therefore hypothesized that upregulation of Na_v_1.1 and Na_v_1.6 subtypes are required for EVS-mediated increases in Na^+^ current amplitude and intrinsic excitability. Knockout of Na_v_1.1, Na_v_1.2 or Na_v_1.6 channels is lethal, resulting in prenatal death in rodent models (Catterall et al., 2010). While heterozygous loss of function of individual Na^+^ channel subtypes causes compensatory changes that often leads to neuronal and circuit hyperexcitability and seizures (Meisler et al., 2021). Therefore, to determine the contribution of individual Na^+^ channel subtypes to homeostatic changes in Na^+^ currents and intrinsic excitability, we used antisense morpholino RNA technology to prevent the upregulation of expression of individual Na^+^ channel subtypes, and then examined the requirement of each Na^+^ channel subtype for EVS-mediated homeostatic increases in Na^+^ current amplitude and intrinsic excitability.

We bulk electroporated the tectum of stage 49 tadpoles with lissamine-tagged, translation-blocking antisense morpholino oligonucleotides (MO) specific for Na_v_1.1, Na_v_1.2, Na_v_1.6, or a control MO. After 24rs, we performed whole cell recordings from lissamine-positive tectal neurons and measured Na^+^ currents from tadpoles exposed to 4hrs of EVS or non-exposed control tadpoles (Figure 5A). We found that upregulation of Na_v_1.6 expression is required for the EVS-mediated increase in the fast Na^+^ current, persistent Na^+^ current and resurgent Na^+^ current; while upregulation of Na_v_1.1 expression is required for the EVS-mediated increase in the persistent and resurgent Na^+^ currents (Figure 5D-F and Figure 5 – figure supplement 2). Additionally, we observed that knockdown of Na_v_1.6 triggers a compensatory increase in the fast Na^+^ current compared to control MO (Figure 5D; *p*=0.0023), providing further evidence for a crucial role for Na_v_1.6 channels in regulating excitability of tectal neurons. In contrast, we observed that upregulation of Na_v_1.2 expression was not required for EVS-mediated increase in the fast, persistent or resurgent Na^+^ currents (Figure 5D-F and Figure 5 – figure supplement 2). Taken together, these data suggest that increased expression of Na_v_1.1 and Na_v_1.6 is the molecular mechanism leading to for EVS-mediated increase in sodium currents.

Our data indicates that the upregulation of Na_v_1.6 expression is required for EVS-mediated increases in sodium currents, and that the specific Na_v_1.6 channel blocker MV1312 decreases intrinsic excitability, suggesting that Na_v_1.6 is a key regulator of sodium currents that control homeostatic changes in intrinsic excitability. Therefore, we next tested whether Na_v_1.6 expression is required for the EVS-mediated homeostatic increase in intrinsic excitability. We found that upregulation of Na_v_1.6 expression is required for the EVS-mediated increases in intrinsic excitability (Figure 5G-H and Figure 5 – figure supplement 3), with upregulation of Na_v_1.6 expression also required for EVS mediated increases in spike amplitude and fast Na^+^ currents (Figure 5I-J and Figure 5 – figure supplement 3). Similar to what was observed when measuring Na^+^ currents, we observed that knockdown of Na_v_1.6 triggers a compensatory increase in the fast Na^+^ current compared to control MO (Figure 5H; *p*=0.0047). As expected, there was no effect of EVS or Na_v_1.6 MO on the amplitude of K^+^ currents (Figure 5K and Figure 5 – figure supplement 3), providing further evidence that regulation of Na^+^ currents is key to the control of intrinsic excitability in tectal neurons. Taken together, these data provide strong evidence that regulation of Na_v_1.6 and Na_v_1.1 mediated Na^+^ currents are key for the homeostatic regulation of intrinsic excitability in *Xenopus* tectal neurons.

### Perturbing expression of Na_v_1.1 and Na_v_1.6 channels during tectal circuit development causes deficits in sensorimotor behaviours

What are the functional consequences of Na^+^ channel subunit mediated regulation of intrinsic excitability during development? The optic tectum, the primary visual area in the tadpole brain, undergoes activity-dependent refinement during development, which is necessary for the accurate performance of visually-guided behaviours (Dong and Aizenman, 2012; Khakhalin et al., 2014; Shen et al., 2014; James et al., 2015; Hamodi et al., 2016). Activity-dependent refinement of tectal circuitry from developmental stage 46 leads to better visual acuity as receptive field size decreases, and the temporal window for multisensory integration becomes narrower, while interventions that alter this activity-dependent process cause perturbed performance in tests of visual acuity, multisensory integration and schooling behaviours (Schwartz et al., 2011; James et al., 2015; Truszkowski et al., 2016). One hypothesis is that the increase in excitability mediated by elevated Na_v_1.1 and Na_v_1.6 levels during developmental stage 46 is important for creating a permissive environment for activity dependent plasticity required for proper tectal circuit development. Thus, we asked whether perturbing expression of sodium channel genes Na_v_1.1 and Na_v_1.6 during tectal circuit development causes behavioural deficits in tasks that require sensorimotor transformations such as visual acuity, multisensory integration and schooling. For these experiments, tadpoles were electroporated with Na_v_1.1 + Na_v_1.6 MOs, or a control MO, at stage 44-45 to suppress Na^+^ channel expression, with the behavioural tasks performed at stage 49.

Visual acuity behavior correlates with the capacity of tectal neurons to tune visual spatial frequency sensitivity (Schwartz et al., 2011), providing a measure of tectal circuitry development. Tadpoles change their swimming speed when presented with counterphasing gratings proportional to the spatial frequency of the gratings (Schwartz et al., 2011). Visual acuity responses were measured by calculating the change in velocity of tadpoles in response to the onset of a series of counterphasing sine wave gratings of different spatial frequencies presented (Figure 6A; 3, 4.5, 9 and 18 cycles/cm). A response was designated if the change in velocity to the stimulus presentation was >150% of the response to no stimulus. Control morphant tadpoles showed increased responsiveness to broad bands with low spatial frequency (Figure 6B), consistent with what has previously described (Schwartz et al., 2011). In contrast, Na_v_ morphant tadpoles showed decreased responsiveness to low spatial frequencies (Figure 6B), indicating a decreased visual acuity. This change in response observed between control MO and Na_v_ morphant tadpoles was not the result of decreases motility, as baseline motility was unchanged between control and Na_v_ morphant tadpoles (Figure 6C). These data suggest that expression of Na_v_1.1 and Na_v_1.6 during a critical period of tectal circuit development is important for the development of visual acuity.

**Figure 6:**
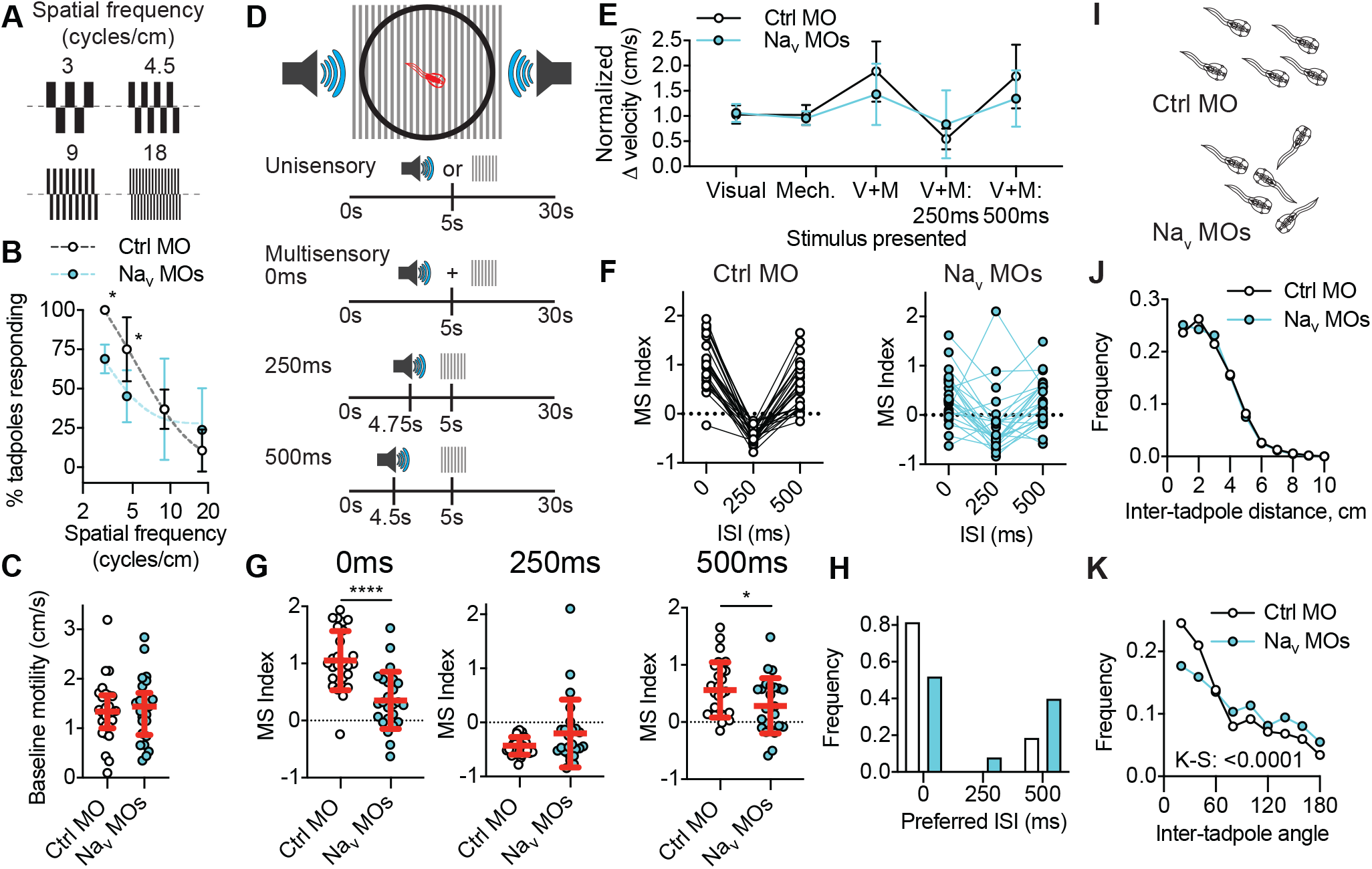
Tadpoles in which expression of Na_v_1.1 and 1.6 was perturbed during tectal circuit development show impairments in visual acuity, multisensory integration and schooling behaviours. Effect of knocking down of Na_v_1.1 and Na_v_1.6 expression at stage 44-46 on tadpole behavior at stage 49. **(A-C)** Effect of developmental knockdown of Na_v_ expression on visual acuity behavior. **(A)** Control morpholino (Ctrl MO) or Na_v_1.1/Na_v_1.6 morpholino (Na_v_ MO) tadpoles were exposed to gratings counterphasing at 4Hz over a range of spatial frequencies (3, 4.5, 9 and 18 cycles/cm). **(B)** Knockdown of Na_v_1.1 and Na_v_1.6 during the critical period of tectal development impaired responses to low spatial frequencies (*N*=7 experiments of 3-4 animals). **(C)** This effect of Na_v_ knockdown is not caused by a change in motility (*n* = 25-27 animals). **(D-H)** Effect of developmental knockdown of Na_v_ expression on multisensory integration behavior. **(D)** Experimental paradigm illustrating the presentation of visual and mechanosensory stimuli, or multisensory stimuli with interstimulus intervals ranging from 0-500ms. **(E)** Mean normalized change in velocity (cm/s) in response to unisensory and multisensory stimuli for Control MO and Na_v_ MO tadpoles. **(F)** Multisensory (MS) indexes calculated for individual tadpoles at 0, 250 and 500ms. Values for individual tadpoles are connected by a line. **(G)** Quantification of MS index for interstimulus intervals of 0, 250 and 500ms. **(H)** Histogram of preferred interstimulus intervals (ISI) for each tadpole. (*N*=8 experiments of 3-4 animals). **(I-K)** Effect of developmental knockdown of Na_v_ expression on schooling behaviour **(I)** Schematic illustrates aggregated schooling behavior observed in control MO and Na_v_ MO tadpoles, with Na_v_ MO tadpoles observed to be less likely to be swimming in the same direction, without a change in inter-tadpole distance. These observations are quantified by observing **(J)** inter-tadpole distance (cm), and **(K)** inter-tadpole angles (*N*=6 experiments of 20 animals per experimental group).

Multisensory integration (MSI), one of the primary functions of the tectum, is a highly conserved property of both neuronal output and behavior whereby the response to a stimulus of a single sensory modality is modulated by the coincident presentation of a stimuli from a different sensory modality. MSI depends on the strength of the neuronal response to each individual unimodal stimulus, the overlap between spatial receptive fields for the two sensory modalities, and the time window between presentation of the cross-modal pair (Wallace et al., 1998; 2006). As tectal neurons mature they become more narrowly tuned to a more diverse range of interstimulus intervals, which occurs congruent with activity-dependent strengthening and refinement of synaptic connections (Felch et al., 2016). Because of its complex nature, MSI is a robust readout of connectivity deficits within the optic tectum. We therefore asked whether perturbing expression of sodium channel genes Na_v_1.1 and Na_v_1.6 during tectal circuit development affects MSI. Subthreshold visual or mechanosensory stimuli, or multisensory stimuli with interstimulus intervals of 0, 250 and 500ms were presented to each tadpole (Figure 6D). Responses were determined by measuring normalized change in velocity of tadpoles to the stimulus onset (Figure 6E), from which we then calculated multisensory (MS) index (Figure 6F). We observed that control tadpoles robustly respond to multisensory stimuli, with tadpoles increasing swimming to interstimulus intervals of 0 or 500ms, and slowing down to 250ms interstimulus intervals (Figure 6G). When we examined the preferred ISI of control tadpoles, we observed that tadpoles most strongly respond to multisensory stimuli with an ISI of 0ms, with a few tadpoles preferring an ISI of 500ms (Figure 6H). In contrast, Na_v_ morphant tadpoles exhibited a less robust response to multisensory stimuli and showed less temporal preference. Na_v_ morphant tadpoles exhibited a decreased MS index for both 0ms and 500ms (Figure 6F-G), with a shift in the preferred ISI that reflected how Na_v_ morphant tadpoles broadly responded to a wider range of ISI (Figure 6H). Taken together, these data show how perturbing Na_v_ expression during a critical period of tectal circuit development affect the development of MSI, consistent with Na^+^ channel expression being important for precise homeostatic regulation of intrinsic excitability.

Tadpoles perform a social aggregation behaviour known as schooling, where tadpoles in close proximity engage in a coordinated unidirectional group swimming, which requires the integration of visual, mechanosensory and olfactory cues. Schooling tadpoles display short inter-tadpole distances and inter-tadpole angles less than 45° with neighboring tadpoles, however, this is altered when tectal circuitry development is perturbed (James et al., 2015; Truszkowski et al., 2016). We had observed that perturbing Na^+^ channel gene expression during tectal circuit maturation causes defects in visual acuity and multisensory integration behaviors; therefore, we predicted that perturbing expression of Na^+^ channel genes Na_v_1.1 and Na_v_1.6 during tectal circuit development would also affect schooling behaviour. Consistent with previous reports, the inter-tadpole distance of control tadpoles was ∼2-3cm, with control tadpoles most likely to be swimming in the same direction as their neighbors (Figure 6J-K). However, when we compared the inter-tadpole distances and angles for Na_v_ morphant tadpoles with control tadpoles, we observed that the inter-tadpole angles of Na_v_ morphant tadpoles was significantly altered, with fewer tadpoles swimming in the same direction (Figure 6K). Whereas there was no effect on the inter-tadpole distance (Figure 6J). These data illustrate how perturbing the homeostatic regulation of intrinsic excitability by modulation of Na^+^ channel gene expression during development alters circuit function and causes impaired social aggregation behaviour, most likely as by perturbing multisensory integration in the tectum.

All of these behaviour tasks require proper activity-dependent refinement of tectal circuits. Tadpoles in which expression of Na_v_1.1 and Na_v_1.6 was perturbed during tectal circuit development show decreased visual acuity, broader multisensory tuning, and abnormal schooling orientation. Hence, these data illustrate the vital importance of the correct regulation of Na^+^ channel gene expression for facilitating normal tectal circuit maturation.

## DISCUSSION

In this study, we sought to determine molecular mechanisms by which neurons of the optic tectum adapt their intrinsic excitability during circuit development and with sensory experience, which has important implications for understanding retinotectal circuit formation. Our data show that neurons of the optic tectum homeostatically adapt their intrinsic excitability during development and in response sensory experience by altering the amplitude of fast, persistent and resurgent Na^+^ currents. Critically, we show that this adaption required changes in expression of sodium channel subtypes Na_v_1.6 and Na_v_1.1, which is necessary for sensory experience-dependent homeostatic increases in Na^+^ current amplitude and intrinsic excitability. Furthermore, we show that this mechanism is critical for the functional development of the retinotectal circuity, as dysregulation of Na_v_1.1 and Na_v_1.6 expression during a key period of development, when the tectal circuitry is undergoing activity-dependent strengthening and refinement, causes deficits in behaviors that depend on visual and multisensory processing. Taken together, these findings illustrate the critical role that dynamic regulation of Na^+^ channel gene expression plays in the homoeostatic regulation of neuronal excitability, and the importance of this process for normal circuit development. The mechanism that we describe improves our understanding of the molecular determinates of excitability in the developing nervous system and highlights the need to better understand the role of Na^+^ channel subtype-specific current adaption in regulating circuit formation during nervous system development.

### A key role for persistent and resurgent Na^+^ currents in regulating homeostatic changes in neuronal excitability

In our search for the cellular mechanisms that regulate homeostatic changes in excitability during retinotectal circuit development, we identified distinct persistent and resurgent Na^+^ currents in neurons of the optic tectum. While persistent currents had previously been identified in these cell types (Aizenman et al., 2003; Hamodi and Pratt, 2014), these had not been characterized, nor their function had previously been explored. Moreover, this is the first report that *Xenopus* tectal neurons express a resurgent Na^+^ current. Persistent and resurgent Na^+^ currents in *Xenopus* tectal neurons broadly resemble those described in mammalian neurons; however, there are some important differences. In mammalian neurons, this persistent Na^+^ current usually represents a small fraction of the fast Na^+^ currents (typically ∼1%), with resurgent Na^+^ current typically ∼5-10% of the fast Na^+^ current amplitude (Lewis and Raman, 2014). In contrast, *Xenopus* tectal neurons have relatively large persistent and resurgent Na^+^ currents. Across all developmental stages studied, the amplitude of the persistent Na^+^ current was ∼40% of fast Na^+^ current, and the amplitude of the resurgent Na^+^ current was ∼35-50%. One interpretation of these data is that fast currents have yet to have fully developed in these relatively immature tectal neurons. Tectal neurons generally have relatively small fast Na^+^ currents in the range of 100-300pA, compared to the nA-sized currents in mammalian neurons. It is possible that as tectal neurons continue to mature, their fast Na^+^ current will increase relative to persistent and resurgent Na^+^ currents. Another interpretation is that this relatively large persistent and resurgent Na^+^ currents allows for rapid adaption of action potential waveform and, therefore, excitability. Propagation of excitatory and inhibitory post-synaptic potentials to the soma causes graded voltage change that, if larger than a certain threshold, leads to the initiation of an action potential at the axon initial segment (Aizenman and Linden, 2000; Cudmore and Turrigiano, 2004; Xu et al., 2005). Hence, the integrative function of a neuron is strongly affected by changes in depolarizing currents, including persistent and resurgent Na^+^ currents, that drive membrane voltage closer to threshold potential (Raman et al., 1997). As tectal neurons develop and exist in an environment where sensory experience is continuing to alter synaptic organization and strength, they require mechanisms to rapidly adapt to high and low levels of synaptic input by changing resting membrane potential as well as AP firing rate, which could be achieved via the regulation of persistent and resurgent Na^+^ currents. Our observation that persistent and resurgent Na^+^ currents are regulated with changes in excitability both across development and in response to patterned visual experience supports this idea.

### Na_v_1.6 is a molecular determinate of neuronal excitability in the developing *Xenopus* visual system

Tectal neurons homeostatically regulate the amplitude of their fast Na^+^ currents in responses to changes in synaptic input caused by experience-dependent developmental changes in circuit architecture (Pratt and Aizenman, 2007; Ciarleglio et al., 2015), short-term changes in visual experience (Aizenman et al., 2003; Ciarleglio et al., 2015), as well as in response to the overexpression of K^+^ channels, such as the shaker-like K_v_1.1 and the delayed-rectifier K_v_2.1 (Pratt and Aizenman, 2007; Dong and Aizenman, 2012). In this study, we extend these findings to show that tectal neurons homeostatically adapt persistent and resurgent Na^+^ currents with changes in neuronal excitability. However, the molecular mechanism by which these Na^+^ currents are adapted in response to changing synaptic inputs, and whether this modulation of Na^+^ current amplitudes is necessary to mediate homeostatic changes in excitability, remained unknown. Using a combination of ion substitution experiments, pharmacology and RNA interference technology, we were able to determine that *Xenopus* tectal neurons express TTX-insensitive persistent and resurgent Na^+^ currents that are largely mediated by Na_v_1.6 channels. This was initially puzzling, because *Xenopus* do not express traditional TTX-resistant Na_v_1.5, Na_v_1.8 or Na_v_1.9 channels. Nevertheless, while the fast Na^+^ current was abolished by TTX (Aizenman et al., 2002), we observed no effect of TTX on the persistent and resurgent Na^+^ currents. It was therefore not clear whether the persistent and resurgent Na^+^ currents observed in this study were carried by a distinct channel type or whether *Xenopus* Na^+^ channels can carry both a TTX-sensitive fast Na^+^ current and TTX-insensitive persistent and resurgent Na^+^ currents. Our ion substitution experiments had shown that these TTX-resistant currents were carried by Na^+^ influx, but the molecular determinate of these TTX-resistant currents remained unknown. TTX-resistant Na^+^ currents have been reported in neurons of other frog species (Campbell, 1992; Kobayashi et al., 1993; 1996), supporting the idea that there may be a TTX-insensitive Na^+^ channel subtype in *Xenopus* tectal neurons. Of the Na^+^ channel subtypes expressed in the *Xenopus* brain (Na_v_1.1, Na_v_1.2 and Na_v_1.6), Na_v_1.6 channels are the most evolutionarily distinct (Zakon, 2012), and have been strongly linked with regulation of persistent and resurgent Na^+^ currents (Raman et al., 1997; Khaliq et al., 2003; Enomoto et al., 2007; Patel et al., 2015). We therefore hypothesized that Na_v_1.6 channels may have a role in regulating persistent and resurgent Na^+^ currents and, hence, homeostatic changes neuronal excitability. This hypothesis was supported by our findings that the specific Na_v_1.6 channel inhibitor MV1312 attenuated persistent and resurgent Na^+^ currents in a use-dependent manner, and that knockdown of Na_v_1.6 attenuated the visual experience-dependent upregulation of fast, persistent and resurgent Na^+^ currents; thus, preventing any homeostatic increase in intrinsic excitability. Moreover, while knockdown of Na_v_1.1 also prevented experience-dependent upregulation of persistent and resurgent Na^+^ currents, expression levels were largely unchanged relative to the change observed for Na_v_1.6, suggesting that Na_v_1.1 plays a supporting role in this process. A specific Na_v_1.1 inhibitor will be required to fully characterize the contribution of these channels to the regulation of Na^+^ currents and excitability in tectal neurons. Nevertheless, these findings have important implications for our understanding of the molecular mechanisms that regulate homeostatic changes in neuronal intrinsic excitability for experience-dependent development of the retinotectal circuitry.

How are Na^+^ channels regulated to mediate homeostatic changes in intrinsic excitability? In flies and rodents, activity-dependent regulation of Na^+^ channel gene expression has been shown to involve the translational regulator Pumilio (Mee et al., 2004; Driscoll et al., 2013; Lin and Baines, 2015). Activity-dependent splicing of the *Drosophila* Na^+^ channel (*DmNa_v_*) modulates the persistent Na^+^ current to regulate neuronal excitability (Muraro et al., 2008; Lin et al., 2009), and can promote seizures (Lin et al., 2012). It is not known whether a similar mechanism to control Na^+^ channel gene expression is present in tectal neurons. To better understand the mechanism by which changes in synaptic input are sensed and translated into a change in Na^+^ current amplitude, and whether this involves the regulation of individual Na^+^ channel subtypes or all Na^+^ channel subtypes, future experiments should explore the molecular mechanisms by which Na^+^ channel gene expression is regulated as tectal neurons undergo homeostatic plasticity of intrinsic excitability. It is especially important to understand how this process might become dysregulated in disease. For instance, increased Na^+^ channel gene expression has been linked to disease progression in epilepsy, with Na_v_1.6 expression found to be increased in layer II neurons of the medial entorhinal cortex in a rodent model of status epilepticus (Hargus et al., 2013), and in hippocampal CA3 neurons in a rodent model of kindling seizures (Blumenfeld et al., 2009). Deciphering the cellular mechanisms by which tectal neurons regulate Na^+^ channel expression and function during retinotectal circuit development will be crucial for understanding how neurons can be flexible and respond to massive changes in circuit and synaptic organization with development or in response to sensory experience, while maintaining the ability to robustly respond to synaptic inputs.

### Functional implications of subtype specific homeostatic plasticity of sodium currents

Early life sensory experience during critical periods of development shapes circuits in the brain (Fox, 1992; Crair and Malenka, 1995; Hensch and Fagiolini, 2005). These critical periods represent time windows when neurons and circuits are particularly sensitive to modification. It is thought that disruption of activity-dependent homeostasis processes during these critical periods may contribute to neurodevelopmental disease such as autism spectrum disorder and epilepsy (Meredith et al., 2012; Doll and Broadie, 2014; Mullins et al., 2016). Indeed, reducing neuronal excitability in the optic tectum during early development by overexpressing the shaker-like K_v_1.1 channel results in tectal neurons with larger visual receptive fields, decreased field sharpness and more persistent recurrent network activity (Dong and Aizenman, 2012), illustrating the importance of the tight control of neuronal excitability for the normal development of circuits. Given their role in action potential generation, Na^+^ channels, particularly Na_v_1.6, are well placed to be key regulators of neuronal excitability. Indeed, both gain-of-function and loss-of-function mutations in Na^+^ channel genes can trigger neuronal and circuit dysfunction that results in neurodevelopmental disorders including epilepsy and autism spectrum disorder (Cannon and Bean, 2010; Meisler et al., 2021). In the mammalian brain, Na_v_1.2 is replaced by Na_v_1.6 as the major constituent of the axon initial segment during postnatal development (Caldwell et al., 2000; Boiko et al., 2001). Neurons of Na_v_1.6-deficient mice found to be have reduced resurgent currents and impaired ability to fire repetitively (Raman et al., 1997; Khaliq et al., 2003; Van Wart and Matthews, 2006), even though there is a compensatory increase in Na_v_1.2 at the axon initial segment and nodes of Ranvier in these Na_v_1.6-deficient neurons (Vega et al., 2008). These data suggest that there may be a subtype specific role of Na_v_1.6 in regulating neuronal intrinsic excitability. Consistent with this idea, we observed that Na_v_1.6 is transiently upregulated at developmental stage 46, which reflects a key period of tectal circuit maturation where activity-dependent refinement and strengthening of synaptic connections. In addition, we found that Na_v_1.6 and the accessory subunit Na_v_4β, which mediate resurgent Na^+^ currents by binding to Na_v_1.6, were highly upregulated in response to sensory experience, and that this was necessary for experience-dependent homeostatic increases in intrinsic excitability. Furthermore, when we prevented upregulation of Na_v_1.6 and Na_v_1.1 expression during the critical period of tectal circuit development at stage 46, we observed deficits in behaviors that rely on sensory integration in the tectum. These findings suggest that perturbing Na^+^ channel mediated regulation of excitability is sufficient to cause deficits in circuit function. By showing the Na^+^ channel subtype specific role in regulating excitability, our work illustrates the utility of the *Xenopus* visual system as an experimental model for studying the effect of Na^+^ channelopathies on neuronal and circuit development.

## MATERIALS AND METHODS

### Animals

All animal experiments were performed in accordance with and approved by Brown University Institutional Animal Care and Use Committee standards. *Xenopus laevis* tadpoles were raised in 10% Steinberg’s solution on a 12hr light/dark cycle at 18-21 °C, with developmental stages determined according to established standards (Nieuwkoop and Faber, 1994). Under these rearing conditions tadpoles generally reach stage 42 at 6-7 days post-fertilization (dpf), stages 44-46 at 9-12 dpf, and stage 49 at 18-20 dpf. All experiments were performed with matched controls from the same clutch of embryos and from at least 3 separate breedings.

### Enhanced visual stimulation (EVS)

To trigger homeostatic increases in the intrinsic excitability of tectal neurons, tadpoles were exposed to short-term (4hr) enhanced visual stimulation at stage 49(Aizenman and Cline, 2007). Freely swimming tadpoles were transferred to a custom built light chamber consisting of four rows of three LEDs that flashed in sequence at 1 Hz to simulate motion stimuli (Sin et al., 2002; Aizenman et al., 2003). Brains were prepared for electrophysiology or qPCR immediately following EVS exposure.

### Morpholinos

Knockdown of expression of Na^+^ channel subtypes was achieved using 3’-lissamine-tagged translation-blocking anti-sense morpholino oligonucleotides (MO; GeneTools, Oregon, USA) targeted to *Xenopus* Na_v_1.1 (5’-TTACTGCTTTGCTACTTTCATAATG-3’), Na_v_1.2 (5’-GTGGTTGCTCCATCTTCTCATCC-3’), and Na_v_1.6 (5’-CAACTTCTCCTGTTAAGTAGCGCCT-3’). To control for off-target effects of MO electroporation, we used a 3’-lissamine-tagged Control MO (5’-CCTCTTACCTCAGTTACAATTTATA-3’). MOs were dissolved in water at 0.1mM. To electroporate MOs into cells of the optic tectum, MOs were injected into the brain ventricle before platinum electrodes were placed on either side of the midbrain and 3-5 pulses at ∼30V with an exponential decay of 70ms applied bidirectionally, which results in bulk electroporation of tectal neurons.

### Electrophysiology

For whole-cell recordings from tectal neurons tadpole brains were prepared as described previously (Wu et al., 1996), to access the ventral surface of the tectum. In brief, tadpoles were anaesthetized in 0.02% tricainemethane sulfonate (MS-222), before brains were filleted along the dorsal midline, removed and pinned to a sylgard block submerged in a recording chamber and maintained at room temperature for the duration of the experiment. Unless otherwise stated, brains were maintained in HEPES-buffered extracellular saline (in mM: 115 NaCl, 2 KCl, 3 CaCl_2_, 3 MgCl_2_, 5 HEPES, 10 glucose, pH 7.2, Osmolarity: 250 mOsm) for the duration of the experiment (typically 2-3hrs). To access tectal neurons the ventricular membrane was removed by suction using a broken glass pipette. Recordings were restricted to the middle one-third of the tectum to avoid any developmental variability along the rostrocaudal axis (Khakhalin and Aizenman, 2012; Hamodi and Pratt, 2014). Cells were visualised using a Nikon eclipse E600FN light microscope equipped with a 60x water-immersion objective, a Lumencor sola light engine for fluorescent illumination, and a Hamamatsu IR-CCD camera. To record from MO electroporated neurons, lissamine-tagged MOs were visualised using fluorescence before switching to an IR-filter for recordings. Whole cell voltage-clamp recordings were performed using glass micropipettes (8-12 MΩ). Electrical signals were measured with a Multiclamp 700B amplifier, digitized at 10 kHz using a Digidata 7550B low-noise analog-to-digital board, and acquired using pClamp 11 software (Molecular Devices). To isolate active currents, leak subtraction was performed in real time using pClamp software. Membrane potential was not adjusted for a predicted 12 mV liquid junction potential. Neurons with series resistance >50 MΩ were not included in the data set. Recordings were analyzed using Axograph X software (John Clements).

To record mixed-current responses, pipettes were filled with K-gluconate intracellular saline solution (in mM: 100 K-gluconate, 8 KCl, 5 NaCl_2_, 1.5 MgCl_2_, 20 HEPES, 10 EGTA, 2 ATP, 0.3 GTP, pH 7.2, Osmolarity: 255 mOsm). To record Na^+^ currents in the absence of K^+^ currents, pipettes were filled with Tris-based intracellular saline solution (in mM: 67 TrisPO_4_, 73 TrisOH, 20 TEA-Cl, 10 EGTA, 10 Sucrose, 2 ATP, 0.1 GTP, pH 7.2, Osmolarity: 255 mOsm). To characterise the ionic composition of currents, we performed ion substitution experiments or used specific blockers in combination with Tris-based internal saline. To abolish all Na^+^ currents in the absence of K^+^ currents, a NMDG-based extracellular saline solution was used (115 NMDG, 2 KCl, 5 HEPES, 10 glucose, 3 CaCl_2_, 3 MgCl_2_, pH 7.2, Osmolarity: 255 mOsm). Fast Na^+^ currents were blocked by the addition of 1-30µM tetrodotoxin (TTX; Tocris Biosciences), Na_v_1.6 currents were inhibited by the addition of MV1312 (5µM; (Weuring et al., 2020)). Na^+^ currents were also targeted with 0.5µM Riluzole (Tocris Biosciences) and 1µM GS967 (Alomone labs). Ca^2+^ currents were blocked by the addition of 100µM CdCl_2_. Ca^2+^ currents were also targeted with 0.5µM ω-Conotoxin MVIIC (CTX; Alomone labs) and 10µM Nimodipine (Tocris Biosciences). To block inward flux of K^+^ ions, we used a TEA-containing external saline solution (in mM: 65 NaCl_2_, 4 KCl, 5 HEPES, 0.01 Glycine, 10 Glucose, 50 TEA-Cl, 3 CaCl_2_, 3 MgCl_2_, pH 7.2, Osmolarity: 250 mOsm).

### Quantification of Na^+^ channel subtype expression levels

To measure Na^+^ channel subtype expression levels during development, whole tecta were harvested from tadpoles at developmental stages 42, 46 or 49. To determine how Na^+^ channel subtype expression was affected by exposure to EVS, whole tecta were harvested from stage 49 tadpoles exposed to 4hr EVS or untreated naïve controls. Optic tecta were collected from 10 tadpoles from 3 independent breedings and stored in RNAlater prior to the isolation of RNA by TRIzol extraction. SuperScript IV VILO first strand cDNA synthesis was then performed according to the manufacturer’s protocol (Invitrogen). Real-time quantitative PCR (RT-qPCR) was performed by using Power SYBR Green master kit with an Applied Biosystems StepOnePlus according to standard manufacturer’s protocols (ThermoFisher). Data were analyzed by the ΔΔCT relative quantification method using the housekeeper gene RSP13, and are represented as a fold change in expression from a control condition (developmental stage 42 or naïve stage 49 control). Primer sequences and corresponding genes are shown in Supplementary Table 1. The RSP13 primer sequences have previously been published (Thompson and Cline, 2016).

### Behaviour experiments

Previous research has demonstrated the behavioural response of *Xenopus* tadpoles to visual and multisensory stimuli, as well as social aggregation behaviour known as schooling. For these experiments, tadpoles were electroporated with Na_v_1.1 + Na_v_1.6 MOs or Control MO at stage 44-45 to perturb Na^+^ channels when intrinsic excitability is highest during development (Pratt and Aizenman, 2007). Behaviour tasks were then carried out at stage 49, with experiments performed in the first 5hrs of the light cycle when tadpoles are most active.

#### Visual acuity

Individual tadpoles were placed in a 5cm diameter dish filled with Steinberg’s solution and exposed to a series of 32 block randomized stimulus presentations of counterphasing sine wave gratings of different spatial frequencies (3, 4.5, 9 and 18 cycles/cm) presented at 4Hz to the tadpoles with a 30s inter-stimulus interval. Gratings were grey scale bars at 80% contrast, which have previously been shown to trigger robust escape responses (Schwartz et al., 2011; Truszkowski et al., 2017). The entire trial lasted approximately 16min. To track tadpoles, the dish was illuminated by 4 IR lights placed uniformly to encircle the dish, with the trial recorded at 30 frames/sec with a SCB-2001 HAD CCD camera (Samsung). Tracking was performed in real time using EthoVision XT (Noldus Information Technology). Tadpoles that did not move over a period of three consecutive minutes were eliminated from the analysis. For analysis, tadpole velocity was averaged over the 1sec pre-stimulus and during the stimulus from which the absolute value of the percent change in velocity was calculated. Values shown are normalized to the change in velocity to no stimulus. A response designated if the change in velocity was >150% of the response to no stimulus. For each experiment, we calculated the % of tadpoles that responded to each spatial frequency. The presentation of counterphasing sine wave gratings were controlled by a MATLAB script to ensure precise timing. Data shown represent individual trials from 27 Ctrl MO and 26 Na_v_1.1+Na_v_1.6 MOs tadpoles from 7 independent experiments. 3 Ctrl MO and 4 Na_v_1.1+Na_v_1.6 MOs tadpoles were excluded from analysis due to inactivity. Data were compared using a Two-way ANOVA with a Holm-Sidak correction for multiple comparisons.

#### Multisensory integration

Individual stage 49 tadpoles were placed in a 5cm diameter dish filled with Steinberg’s solution and exposed to a series of 40 block randomized stimulus presentations of visual, mechanosensory, or visual plus mechanosensory stimuli with inter-stimulus intervals of 0, 250, and 500ms. Stimuli were presented for 2 seconds with an interstimulus interval of 30s, with the entire trial lasting approximately 16min. Visual stimuli consisted of greyscale stripes of 25% contrast that alternated at 4Hz on a CRT monitor located beneath the dish. This stimulus has been shown to be sub-threshold for tadpoles (Truszkowski et al., 2017). That is, visual stimulus alone does not trigger a startle response in tadpoles. Mechanosensory stimuli were low-volume clicks played through two speakers connected to the dish by wooden rods so that they vibrated the liquid of the dish, which activates the ear, skin and lateral line sensory organs. This low-volume stimulus has previously been shown to be sub-threshold using a pre-pulse inhibition protocol (James et al., 2015). For multisensory stimuli, visual stimuli were preceded by mechanosensory stimuli at 500ms, 250ms and 0ms interstimulus intervals. Visual and mechanosensory stimuli were controlled by a MATLAB script to ensure precise timing. To track tadpoles the dish was illuminated by 4 IR lights placed uniformly to encircle the dish, with the trial recorded with a SCB-2001 HAD CCD camera (Samsung). Tracking was performed in real time using EthoVision XT (Noldus Information Technology). Tadpoles that did not move over a period of three consecutive minutes were eliminated from the analysis. To analyse the response of tadpoles to stimulus presentation, tadpole velocity was averaged over the 1sec pre-stimulus and during the stimulus from which the absolute value of the percent change in velocity was calculated. Values shown are normalized to the change in velocity to no stimulus, with all trials from each stimulus condition averaged. From these responses, we then calculated the multisensory (MS) index (equation: [multisensory - unisensory]/unisensory). Data shown was calculated with the visual stimulus used for the unisensory response; however, there was no difference if the mechanosensory stimulus was used. Data shown represent individual trials from 27 Ctrl MO and 25 Na_v_1.1+Na_v_1.6 MOs tadpoles from 7 independent experiments. 3 Ctrl MO and 5 Na_v_1.1+Na_v_1.6 MOs tadpoles were excluded from analysis due to inactivity. Data were compared using a One-way ANOVA with a Holm-Sidak correction for multiple comparisons.

#### Schooling

Schooling experiments were performed as recently described (Lopez et al., 2021), with experiments and analysis conducted using freely available code (https://github.com/khakhalin/Xenopus-Behavior). In brief, Ctrl MO or Na_v_1.1+Na_v_1.6 MOs tadpoles at stage 49 were transferred to a 17cm diameter glass bowl on a LED tracing tablet (Picture/Perfect light pad), which in turn sat atop a dental vibrator (Jintai). Images of the bowl were then captured every 5min over the course of 1hr, with tadpoles dispersed by a vibration that was triggered 150sec prior to each image capture. Images were captured using a GoPro Hero 7 (GoPro Inc.). Vibration delivery and image acquisition were controlled by a custom Python script. For analysis, the position and heading of each tadpole was identified by acquiring the x-y coordinates of the head and gut of each tadpole using the multipoint tool in FIJI. A Delaunay triangulation was then used to calculate inter-tadpole distance and angles between neighboring tadpoles, which were then compared using a Kolmogorov-Smirnov test.

### Statistical analysis

Statistics were performed in Prism 8 (GraphPad Software). Normally distributed data is presented as mean ± SD and analysed using Welch’s *t*-test or Welch’s one-way ANOVA with a Dunnnet T3 test for multiple comparisons. Nonparametric data is presented as median with IQR and analysed with a Mann-Whitney U-test or a Kruskal-Wallis with Dunn’s test for multiple comparisons. Sample sizes were based on power analyses and known biological variability from prior work (Pratt and Aizenman, 2007; Ciarleglio et al., 2015; James et al., 2015; Truszkowski et al., 2017). No outliers were excluded from the data analysis.

## ACKNOWLEDGEMENTS

We thank members of the Aizenman lab for their valuable intellectual input, and especially thank Virgilio Lopez and Mimi Oupravanh for animal care. We are grateful to Dr. Mirko Rivara (Università di Parma, Parma, Italy) for the kind gift of MV1312 that was used in this study. This work was supported by R01 EY027380 awarded to C.D.A from the National Eye Institute.

## AUTHOR CONTRIBUTIONS

**Adrian C Thompson**: Conceptualization, Methodology, Validation, Formal analysis, Investigation, Data curation, Writing - original draft, Writing - review and editing, Visualization. **Carlos D Aizenman**: Conceptualization, Methodology, Writing - original draft, Writing - review and editing, Visualization, Supervision, Project administration, Funding acquisition.

## SUPPLEMENTARY FIGURES

**Figure 1 - figure supplement 1:**
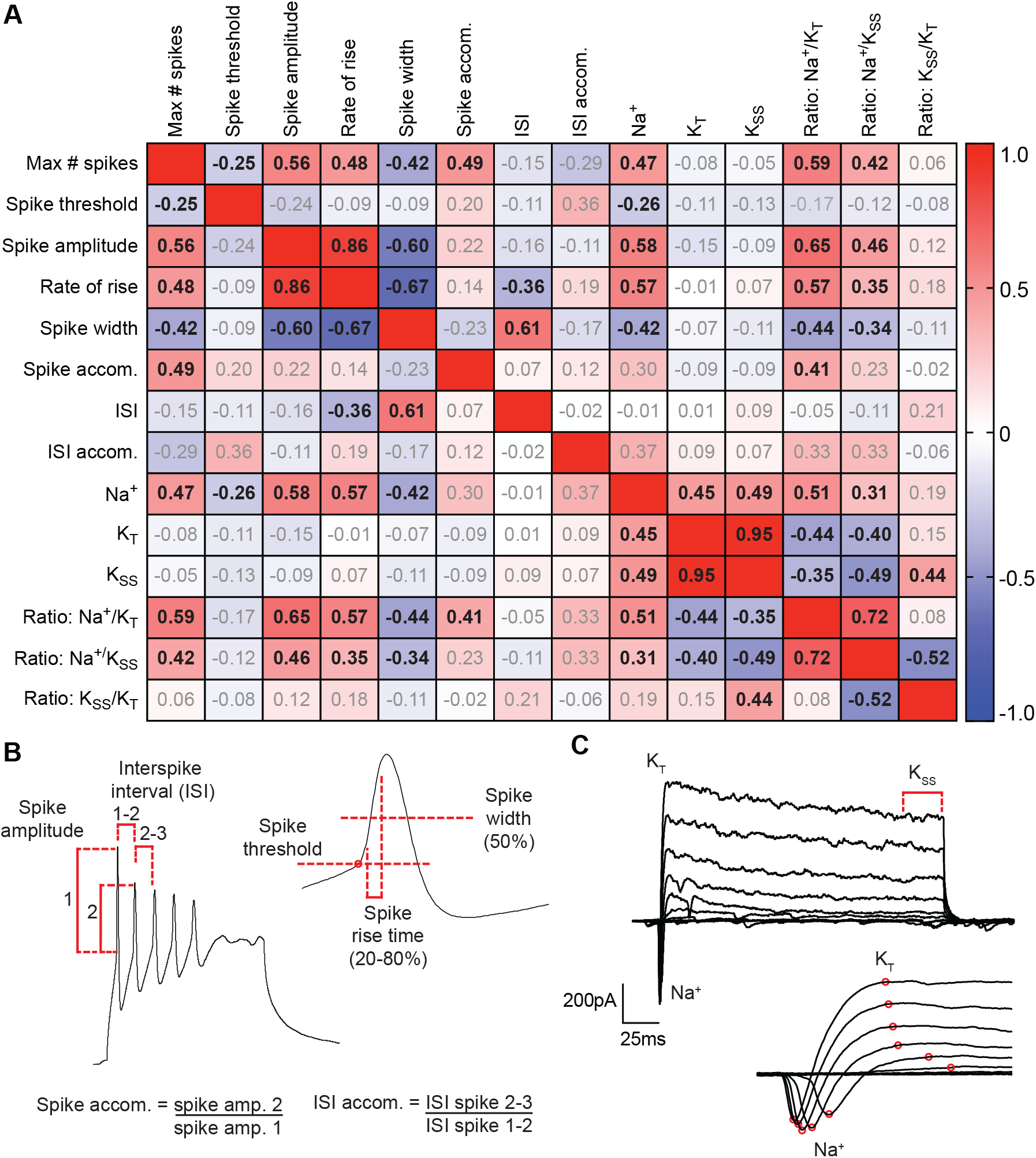
Relationship between biophysical properties of *Xenopus* tectal neurons. **(A)** Pearson correlation matrix showing the relationship in stage 49 tectal neurons between intrinsic excitability and spike characteristics, sodium current (Na^+^), transient potassium current (K_T_), steady state potassium current (K_SS_), and the recovery of Na^+^ currents from fast inactivation. Values represent Pearson correlation values (r), with bolded values indicating significant correlations (*p*<0.05). **(B)** Spiking response to a current step injection. The number of spikes elicited by current step injection was quantified using the following criteria: to qualify as a spike the amplitude of the spike was at least half the amplitude and no wider than three times the width of the first spike. Spike amplitude was measured as the voltage change from threshold voltage to the voltage at spike peak. Spike accommodation was calculated as the amplitude of the 2^nd^ spike divided by the amplitude of the 1^st^ spike. Interspike interval (ISI) was measured as the peak-to-peak time between adjacent spikes. ISI accommodation was calculated as the ISI for spikes 2-3 divided by the ISI for spikes 1-2. *Inset*: An expanded view of the first spike to show the measurements for spike threshold, spike rise time (measured as the time to rise from 20-80% of the spike), and spike width (measured as the width at 50%). **(C)** Mixed current responses to voltage step injections following passive current subtraction. *Inset*: An expanded view of the first 10ms showing the measurement of the temporally distinct Na^+^ and K_T_ currents.

**Figure 2 – Figure supplement 1.**
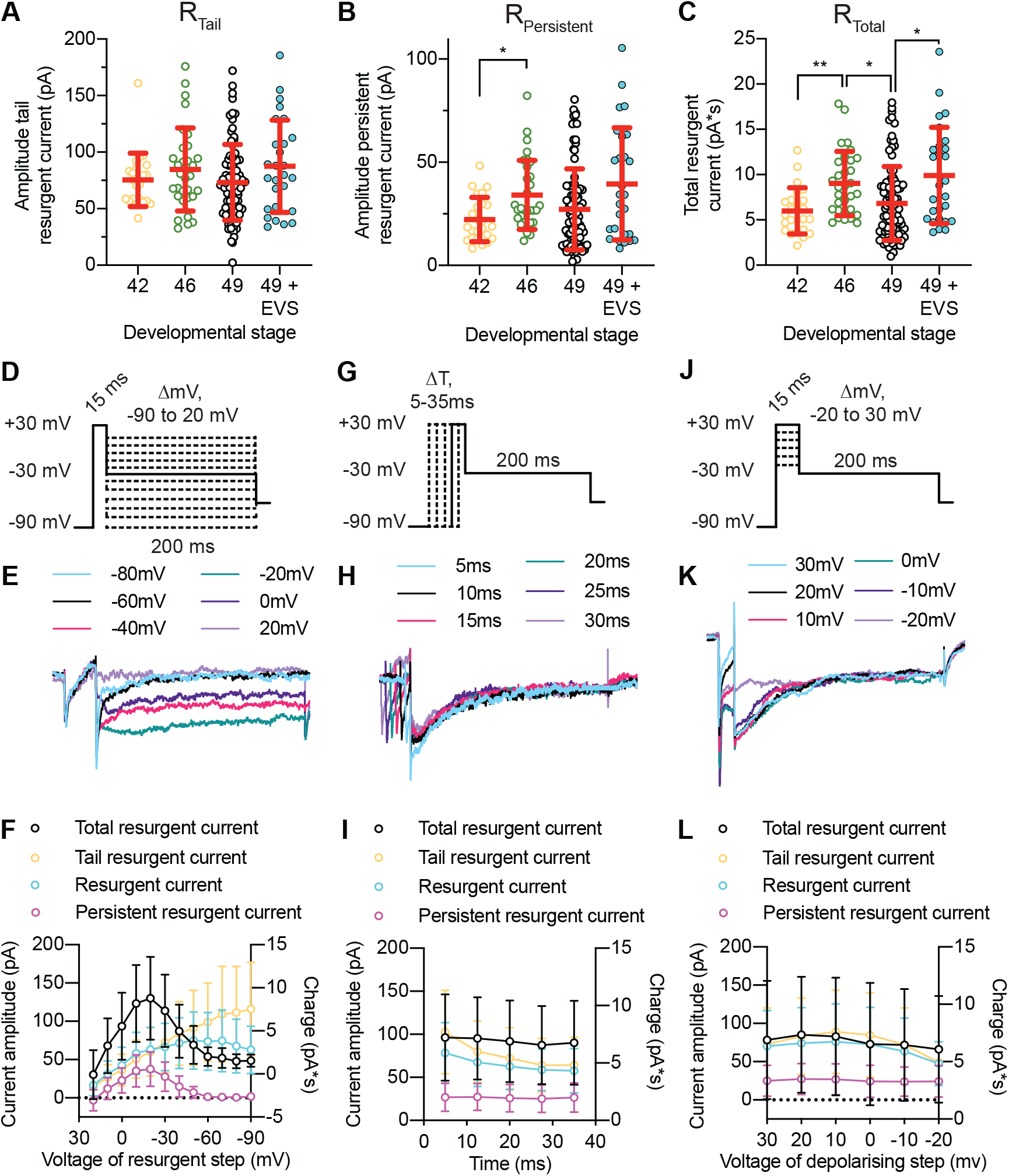
**(A-C)** Quantification of the effect of development and 4hr EVS on **(A)** the tail resurgent Na^+^ current, **(B)** the persistent resurgent Na^+^ current, and **(C)** the total resurgent Na^+^ current. **(D-F)** The effect of altering voltage of the resurgent step from −90mV to +20mV on resurgent Na^+^ currents. **(G-I)** The effect of altering the time of the depolarizing step from 5 to 35ms on resurgent Na^+^ currents. **(J-L)** The effect of altering voltage of the depolarizing step from −20mV to +30mV on resurgent Na^+^ currents.

**Figure 2 – Figure supplement 2:**
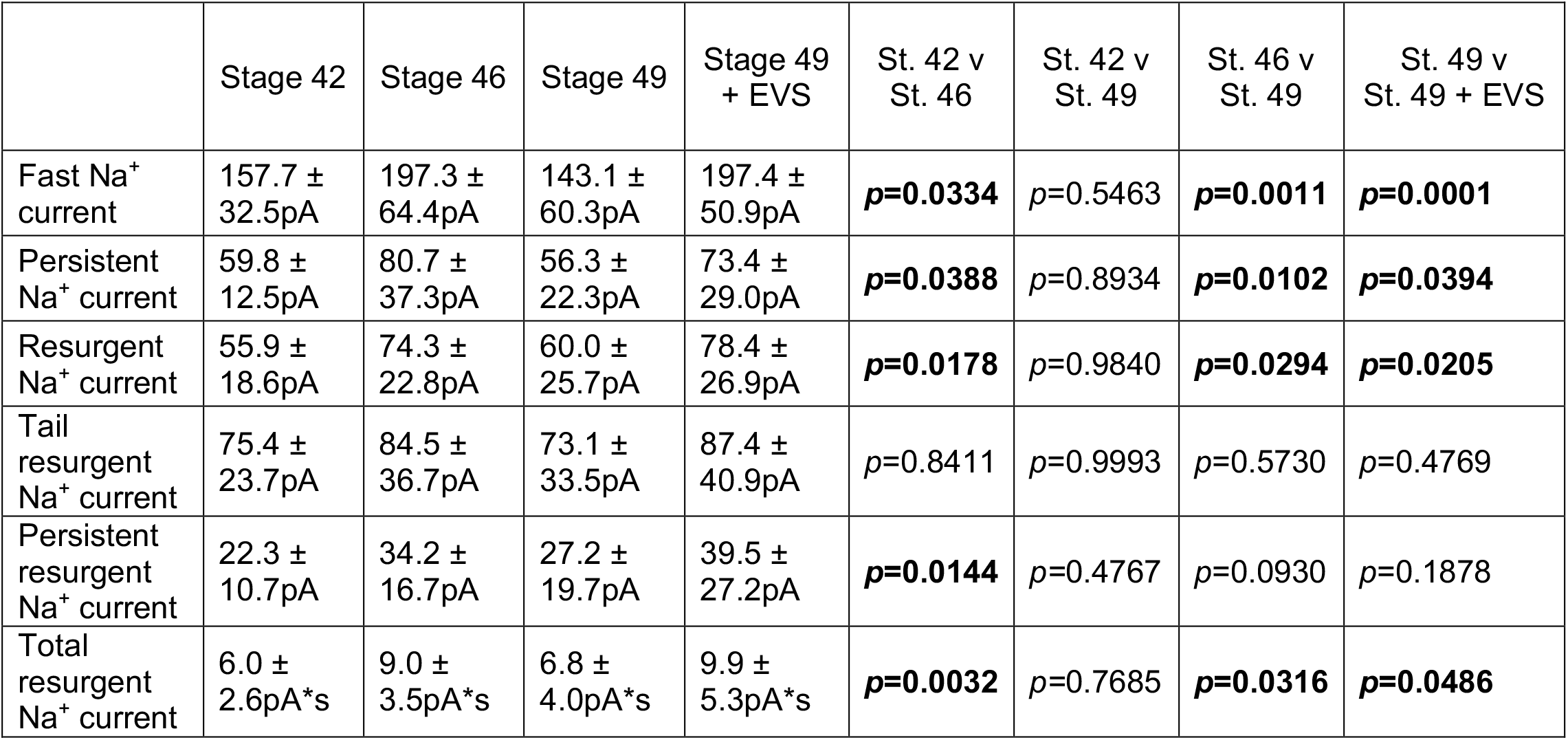
Table shows values (median ± SD) and comparisons between experimental groups for data shown in Figure 2 and Figure 2 – figure supplement 1. Groups were compared using a Welch’s ANOVA test with Dunnett T3 test for multiple comparisons.

**Figure 3 – Figure supplement 1:**
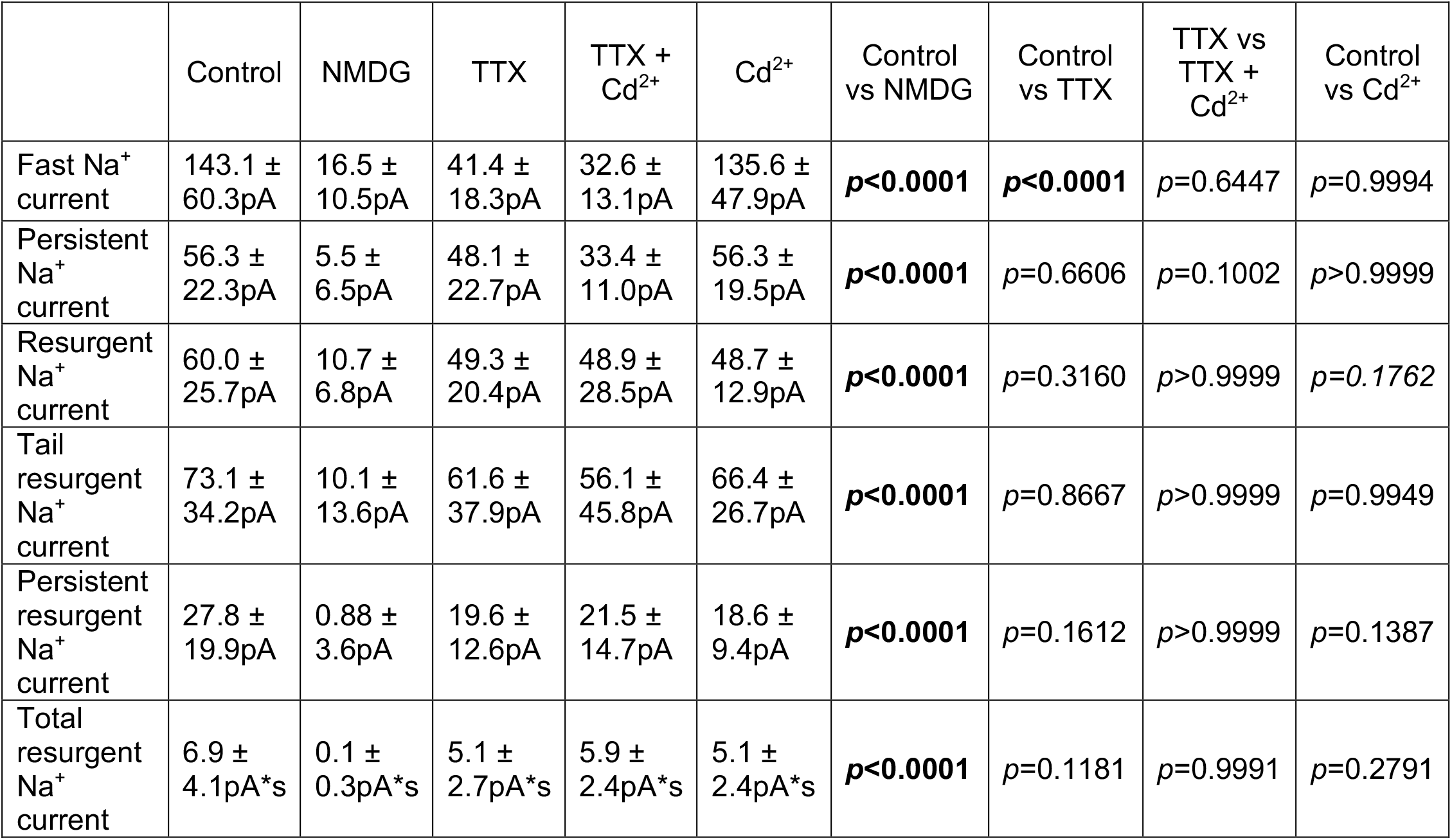
Table shows values (median ± SD) and comparisons between experimental groups for data show in Figure 3. Groups were compared using a Welch’s ANOVA test with Dunnett T3 test for multiple comparisons.

**Figure 3 – Figure supplement 2:**
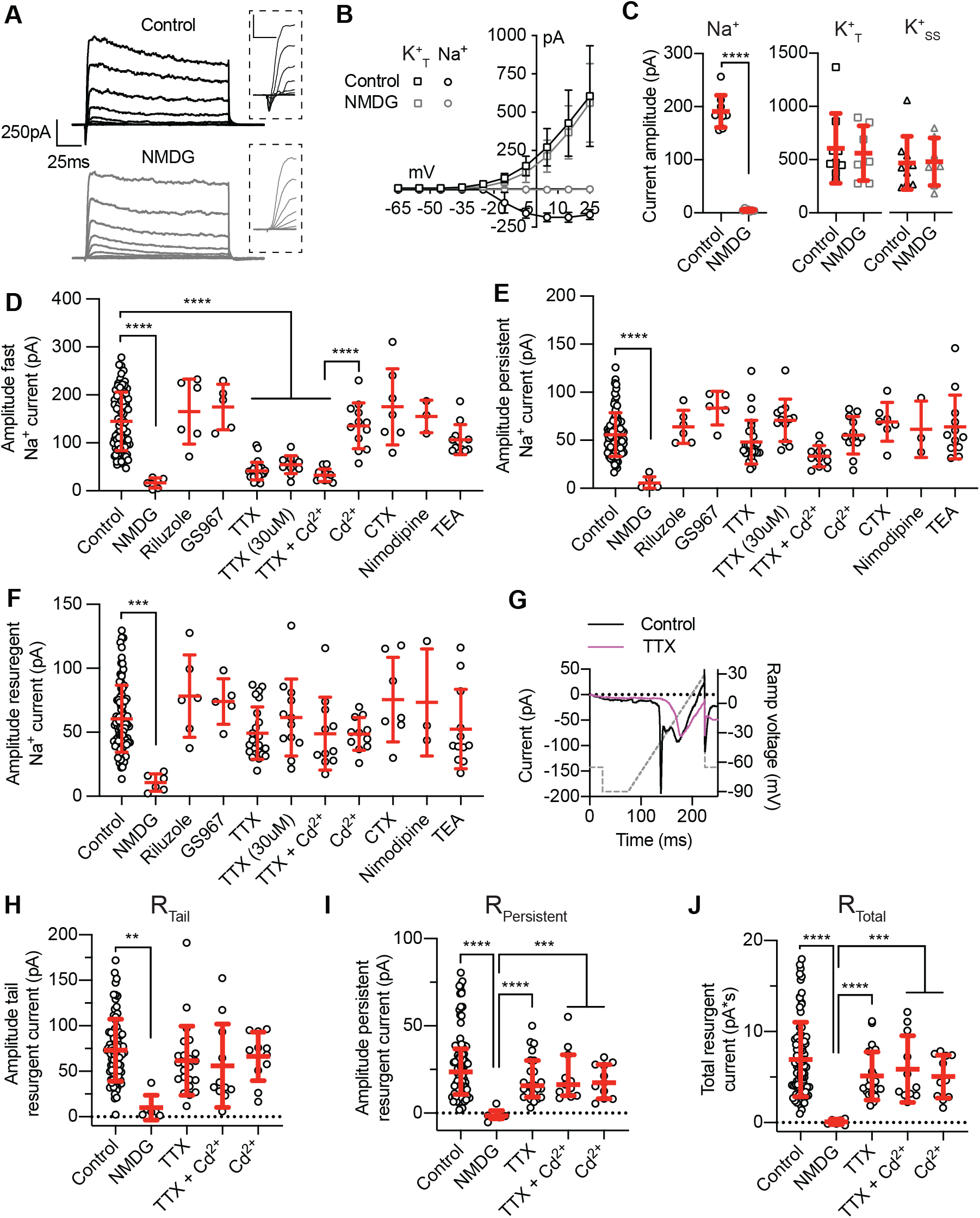
The effect of ion substitution and channel blockers on distinct Na^+^ currents in *Xenopus* tectal neurons. **(A)** Example recordings from stage 49 tectal neurons in normal conditions and following wash in of NMDG-based external saline with zero extracellular Na^+^. *Insets*: Magnification of the initial 7ms of recordings highlighting the effect of NMDG on the fast Na^+^ current. Scale is 50pA and 5ms. **(B)** Averaged current-voltage plots showing the effect of NMDG on fast Na^+^ and peak K^+^ currents. **(C)** Quantification of peak current amplitude for the fast Na^+^ current, the peak K^+^ current, and the steady state current in control and NMDG conditions (\*\*\*\**p*<0.0001; students *t*-test. *n* = 7-9 cells). **(D-F)** Extended plots of those presented in Figure 3 showing the effect of additional ion substitution or specific channel blockers on **(D)** fast Na+ currents, **(E)** persistent Na+ currents, or **(F)** resurgent Na+ currents. **(G)** Plot shows the effect of TTX on the voltage-dependency of the persistent Na^+^ current. Dashed line shows the voltage, black line shows an example control cell, while the magenta line shows a TTX cell. Notice the loss of fast Na^+^ current. **(H-J)** Plots show the effect of NMDG, TTX and Cd on tail and persistent resurgent Na^+^ currents as well as the total resurgent Na^+^ current.

**Figure 4 – figure supplement 1:**
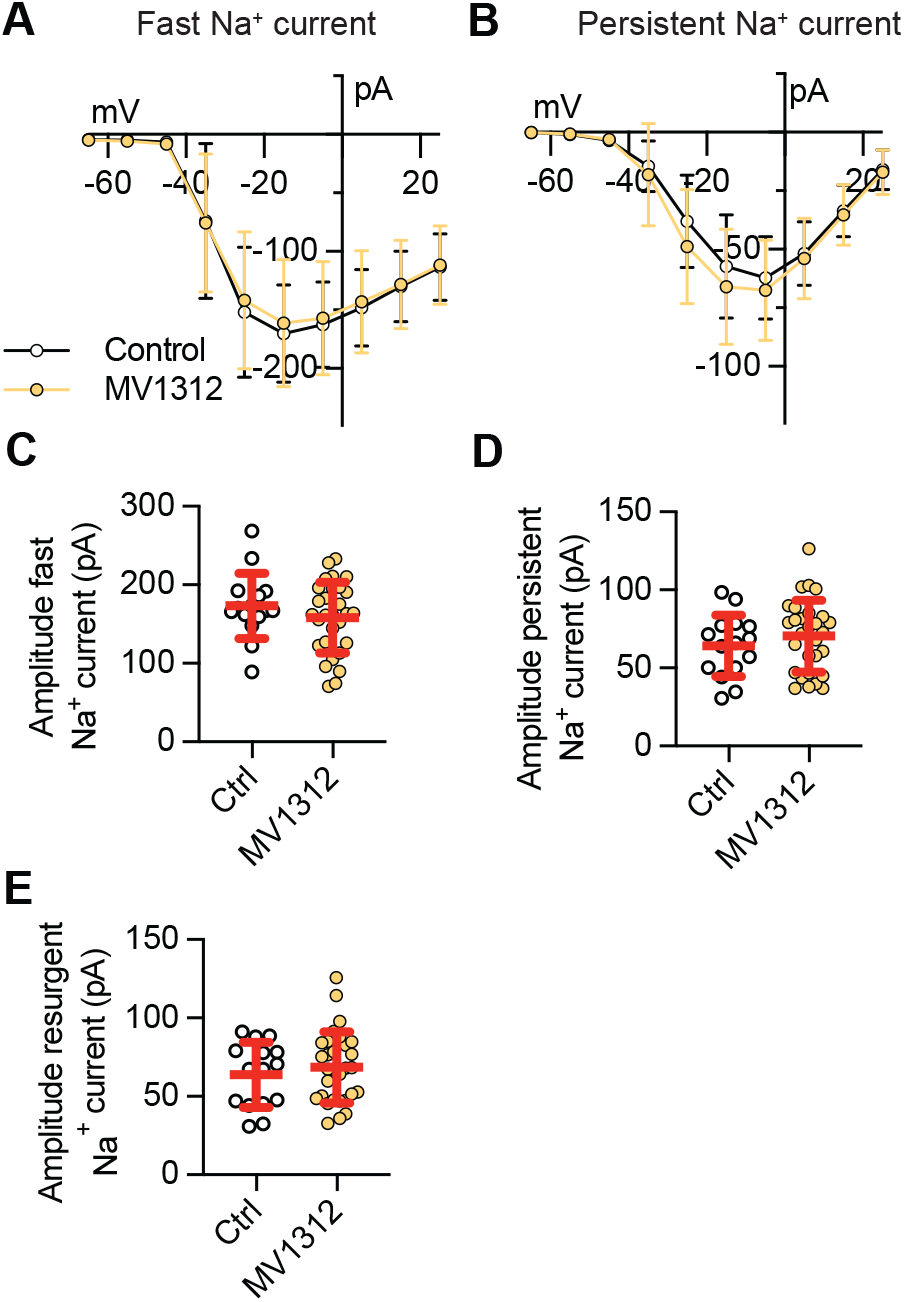
Acute wash in of the Na_v_1.6 specific inhibitor MV1312 has no effect on fast, persistent or resurgent Na^+^ currents. **(A-B)** Averaged *I-V* plots from control stage 49 tectal neurons and stage 49 tectal neurons following wash in of 5µM MV1312. **(C-E)** Quantifications of peak amplitudes for fast, persistent and resurgent Na^+^ currents (median and SD).

**Figure 4 – Figure supplement 2:**
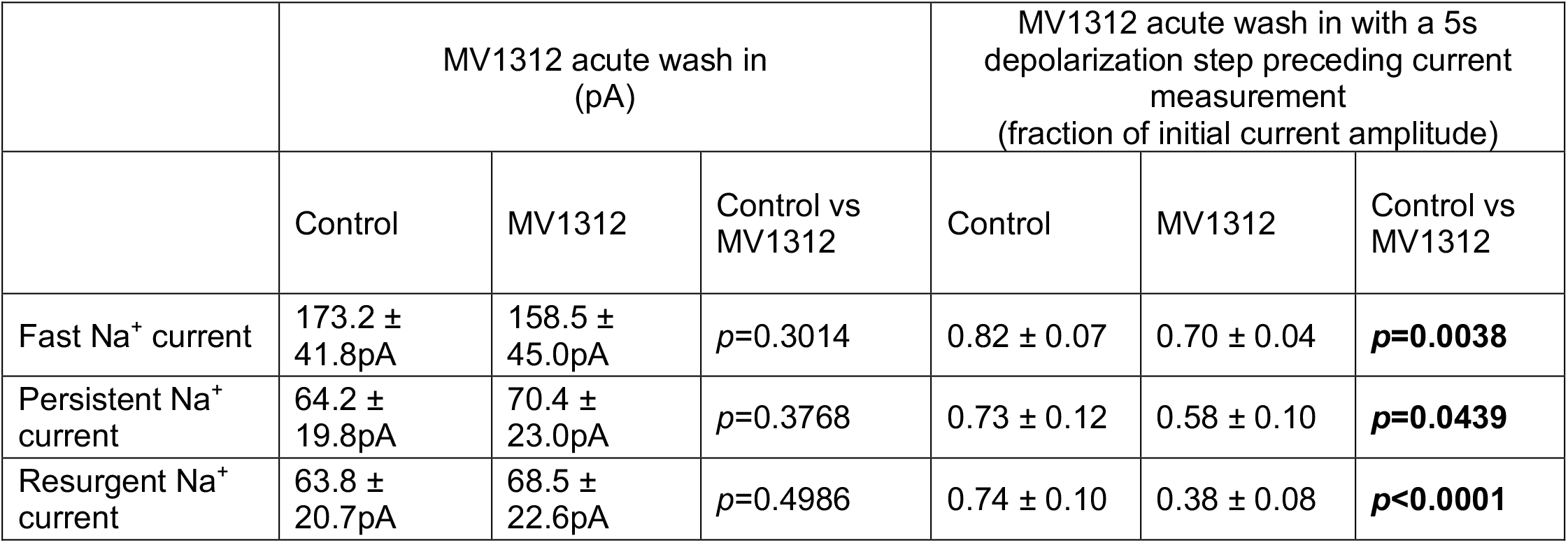
Comparison of fast, persistent and resurgent Na^+^ currents in response to acute wash in of the Na_v_1.6 channel inhibitor MV1312. Table shows values (median ± SD) and comparisons between experimental groups for data show in Figure 4. Wash in (n=15 and 30). Wash in with 5s depolarizing step (n=8 and 5). Groups were compared using a Welch’s ANOVA test with Dunnett T3 test for multiple comparisons.

**Figure 5 - figure supplement 1:**
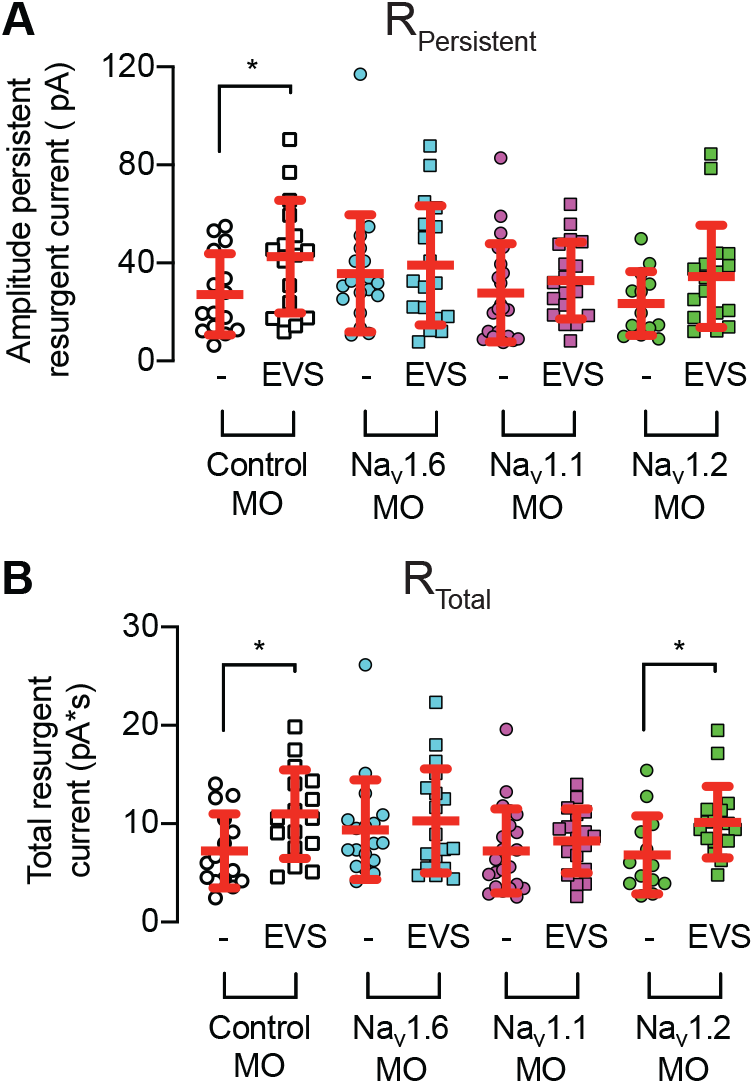
Effect of knocking down expression of Na^+^ channel subtypes on persistent resurgent Na^+^ current and total resurgent Na^+^ current. Exposure of stage 49 tadpoles to 4hrs of EVS triggers an increase in the amplitude of **(A)** persistent and **(B)** total resurgent Na^+^ currents, which is attenuated by the knockdown of Na_v_1.1 and Na_v_1.6.

**Figure 5 - figure supplement 2:**
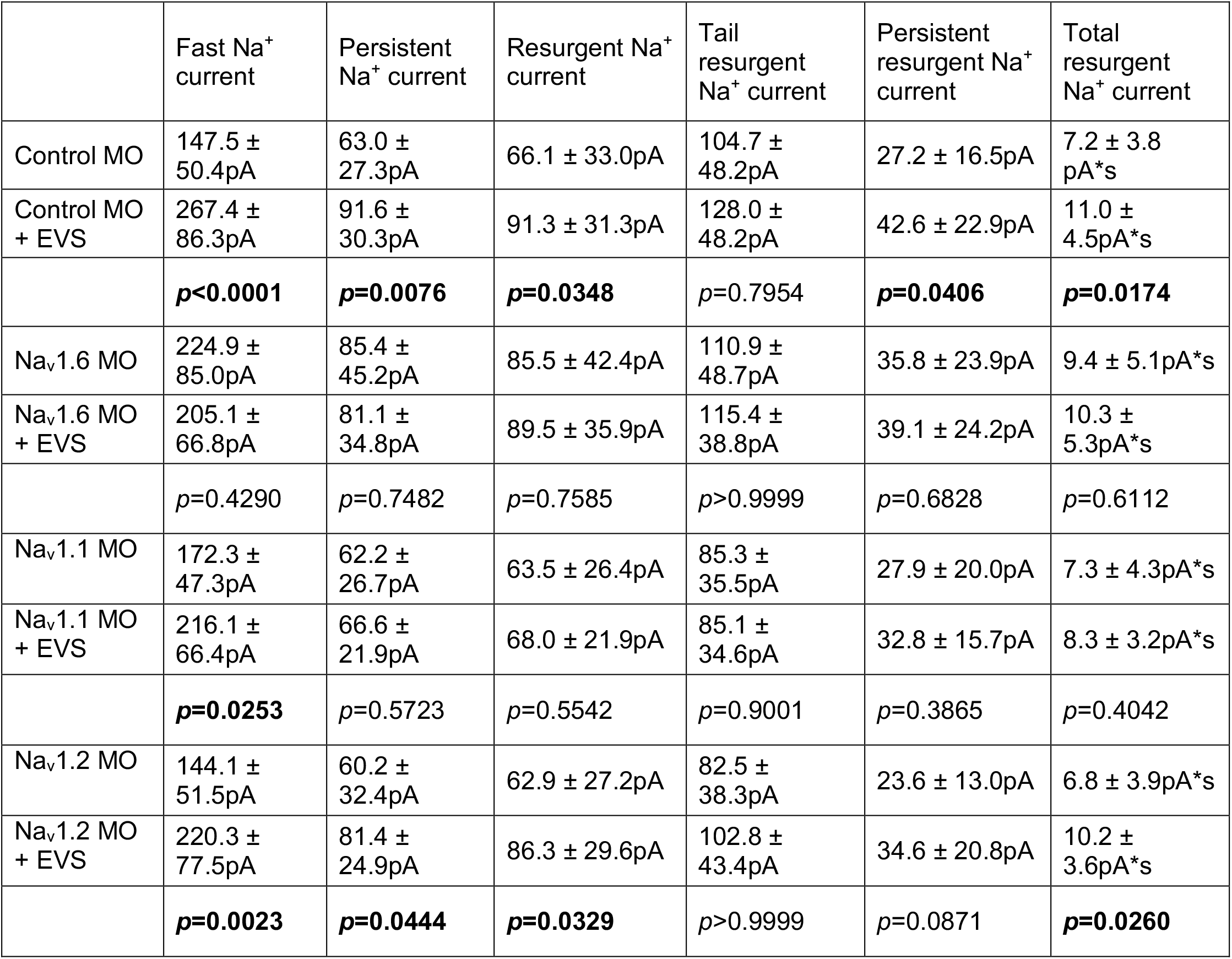
Effect of SCN KD on Na^+^ currents. Table shows values (median ± SD) and comparisons between experimental groups for data show in Figure 5D-F (n=16-22). Groups were compared using a Welch’s ANOVA test with Dunnett T3 test for multiple comparisons.

**Figure 5 - figure supplement 3:**
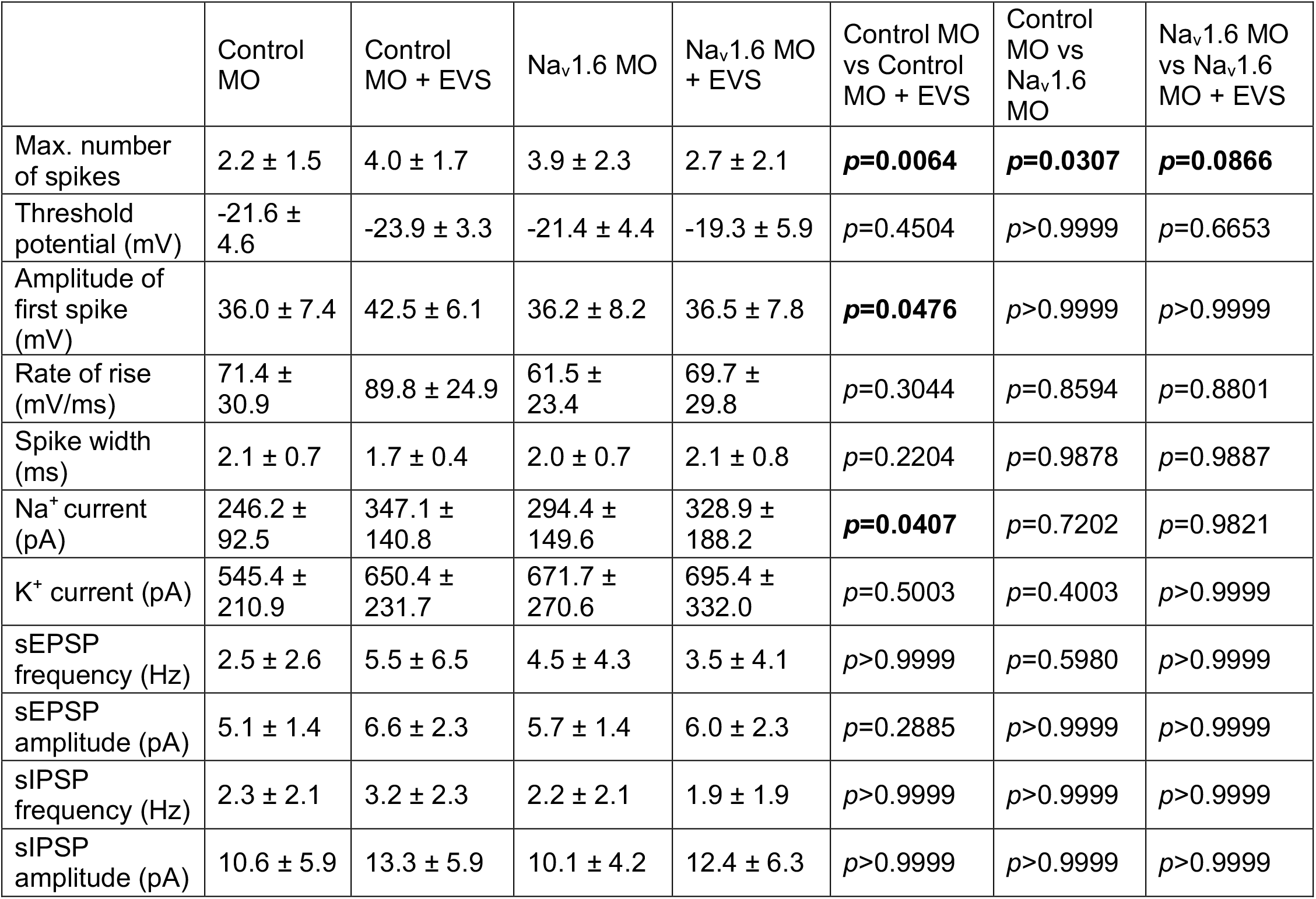
Effect of SCN8A KD on excitability of tectal neurons. Table shows values (median ± SD) and comparisons between experimental groups for data show in Figure 5G-K (n=16-31). Groups were compared using a Welch’s ANOVA test with Dunnett T3 test for multiple comparisons.

**Supplementary Table 1:**
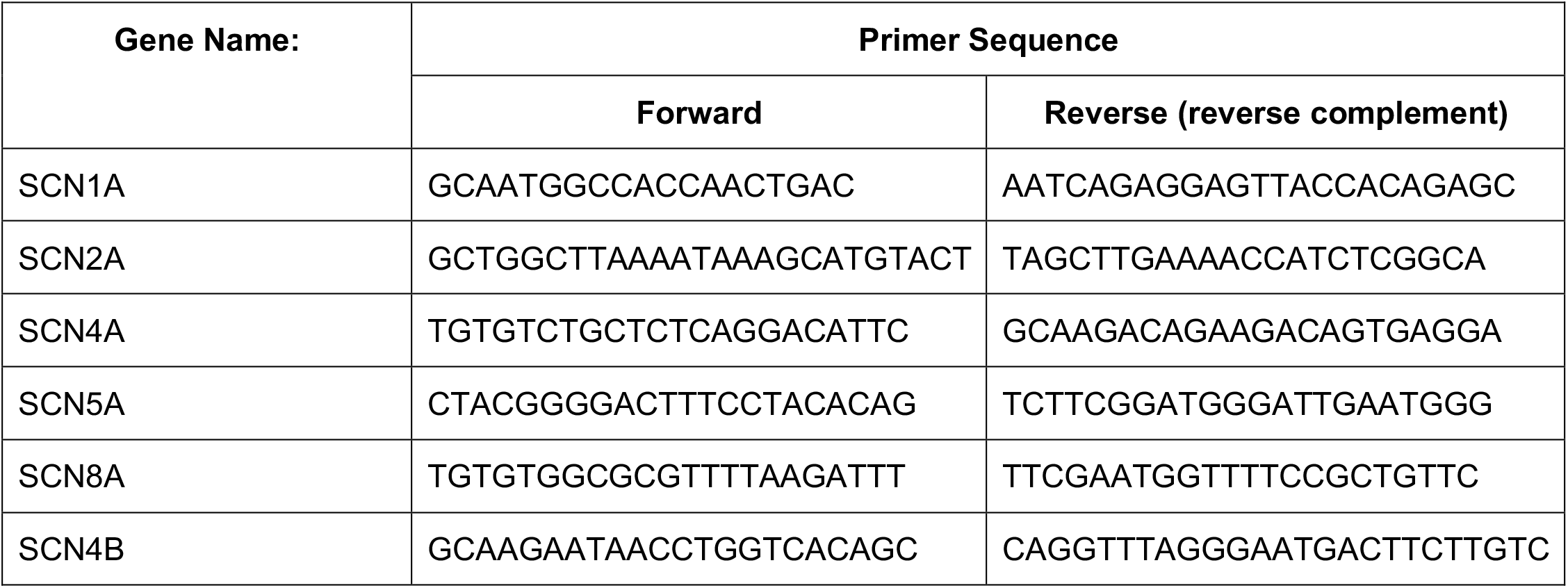
Primer sequences for RT-qPCR experiments targeting alpha and beta subunit genes to determine Na^+^ channel gene expression.

